# The *Aspergillus fumigatus maiA* gene contributes to cell wall homeostasis and fungal virulence

**DOI:** 10.1101/2023.10.18.562787

**Authors:** X Guruceaga, U Perez-Cuesta, A Martin-Vicente, E Pelegri-Martinez, H Thorn, S Cendon-Sanchez, J Xie, A Nywening, A Ramirez-Garcia, JR Fortwendel, A Rementeria

## Abstract

In this study, two distinct *in vitro* infection models of *Aspergillus fumigatus*, using murine macrophages (RAW264.7) and human lung epithelial cells (A549), were employed to identify the genes important for fungal adaptation during infection. Transcriptomic analyses of co-incubated *Aspergillus* uncovered 140 fungal genes up-regulated in common between both models that, when compared with a previously published *in vivo* transcriptomic study, allowed the identification of 13 genes consistently up-regulated in all three infection conditions. Among them, the *maiA* gene, responsible for a critical step in the L-phenylalanine degradation pathway, was identified. Disruption of *maiA* resulted in a mutant strain unable to complete the Phe degradation pathway, leading to an excessive production of pyomelanin when this amino acid served as the sole carbon source. Moreover, the disruption mutant exhibited noticeable cell wall abnormalities, with reduced levels of β-glucans within the cell wall. the *maiA-1* mutant strain induced reduced inflammation in primary macrophages and displayed significantly lower virulence in a neutropenic mouse model of infection. This is the first study linking the *A. fumigatus maiA* gene to fungal cell wall homeostasis and virulence.

## Introduction

Melanins are secondary metabolites composed of indolic monomers or phenolic polymers that act as pigments (1). The saprophytic fungus *Aspergillus fumigatus* can produce two types of melanins. One of them is the dihydroxynaphthalene melanin, also known as DHN-melanin, which is one of the principal components of the conidial cell wall (2) together with galactomannans, β-1,3-glucans, chitin and rodlet proteins. In addition, this fungus can produce a second type of melanin, pyomelanin which is an extracellular water-soluble pigment (3). This molecule is produced by spontaneous polymerization of homogentisate (HGA) during the L-phenylalanine/L-tyrosine (Phe/Tyr) degradation pathway. In this pathway, HGA is accumulated and converted to benzoquinone acetate by spontaneous oxidative transformation, then polymerized to pyomelanin.

Among the biological functions that pyomelanin fulfills, this molecule seems to protect young hyphae against oxidative stress (4). Pyomelanin production is associated with conidial germination and directly depends on three surface sensors (Wsc1, Wsc3 and MidA) that detect Phe or Tyr and send a signal through Rho GTPase (Rho1) and MAP kinases (Bck1, Mkk2 and MpkA) in order to produce the activation of the cluster of genes involved in the Phe/Tyr degradation pathway (5). Furthermore, there are studies that demonstrate an overexpression of the cluster of genes involved in the Tyr degradation pathway when the fungus is exposed to both mechanical and chemical cell wall stresses (5, 6). Therefore, pyomelanin could be a cell wall protective factor in these conditions.

It has also been described that the *A. fumigatus* Phe/Tyr degradation pathway is mediated by six genes located on chromosome 2 (*hppD*, *hmgX*, *hmgA*, *fahA*, *maiA* and *hmgR*). In addition, there are two other genes not included inside the abovementioned cluster, *phhA* and *tat*, which code for phenylalanine hydroxylase and the tyrosine aminotransferase, respectively (7). They oversee the first degradation steps of Phe and Tyr to 4-hydroxyphenylpyruvate. There are several published studies uncovering the function of the genes of the Phe/Tyr degradation gene cluster and their importance for the fungus based on deletion/disruption mutants. However, there are still two genes of the cluster that are not well-characterized because they have not been silenced yet (*fahA* and *maiA*).

The *A. fumigatus maiA* is an exonic 696 bp gene (Afu2g04240) that does not contain introns in its sequence and codifies a maleylacetoacetate isomerase putatively involved in this Phe/Tyr degradation pathway. Specifically, this enzyme orchestrates the isomerization of 4-maleylacetoacetate (4-MA) into 4-fumarylacetoacetate (4-FA) (3). In theory, the resulting phenotype of the defective *maiA* strain should phenocopy Δ*hmgA* (defective *hmgA* strain), which is the gene that codifies the enzyme located just upstream of *maiA* in this degradation pathway, since no other function for this *maiA* gene is known at this time.

In this study, the *A. fumigatus* genes most strongly over-expressed upon co-culture with two different cell lines (murine macrophages and human lung epithelial cells) were studied using the AWAFUGE microarray. Among them, *maiA* was selected because it was overexpressed in these two *in vitro* models, as well as in a previously published *in vivo* infection (8). Therefore, to study its role in fungal biology and virulence, a disruption mutant strain (*maiA-1*) was generated and characterized. The data obtained strongly support the involvement of *maiA* in the maintenance of cell wall structure as well as in the virulence of the fungus using a neutropenic mouse model.

## Results

### Study of co-incubation between *A. fumigatus* and macrophages

The results obtained from co-incubation with the murine macrophage RAW 264.7 cell line revealed a maximum of 80% phagocytosis after 4 hours of co-incubation (Fig. 1A). Progressive production of reactive oxygen species (ROS) by macrophages was also observed, reaching a maximum after 8 hours of contact with the conidia (47.88% more production than macrophages not inoculated) (Fig. 1B). A fast reactive nitrogen species (RNS) production was also detected, with the opposite pattern of ROS, reaching the nitrite maximum production after only 5 minutes (71.93 µM), and decreasing almost completely after 80 minutes of contact (Fig. 1C). As expected, these activities affected the viability of the conidia, which decreased progressively over the time of contact with macrophages (Fig. 1D). However, the surviving conidia seem to adapt to the stress caused by the interaction with the phagocytes. In fact, although the germination of conidia reached 30% at 6 hours in all cases (infection and control cultures), at the end of the experiment significantly higher germination was detected in the conidia incubated with macrophages, compared to the controls (Fig. 1E). In contrast, we did not find any differences in the amount of hyphal growth or their ability to stablish a second axis of polarity during germination (Fig. 1F).

**Figure 1.**
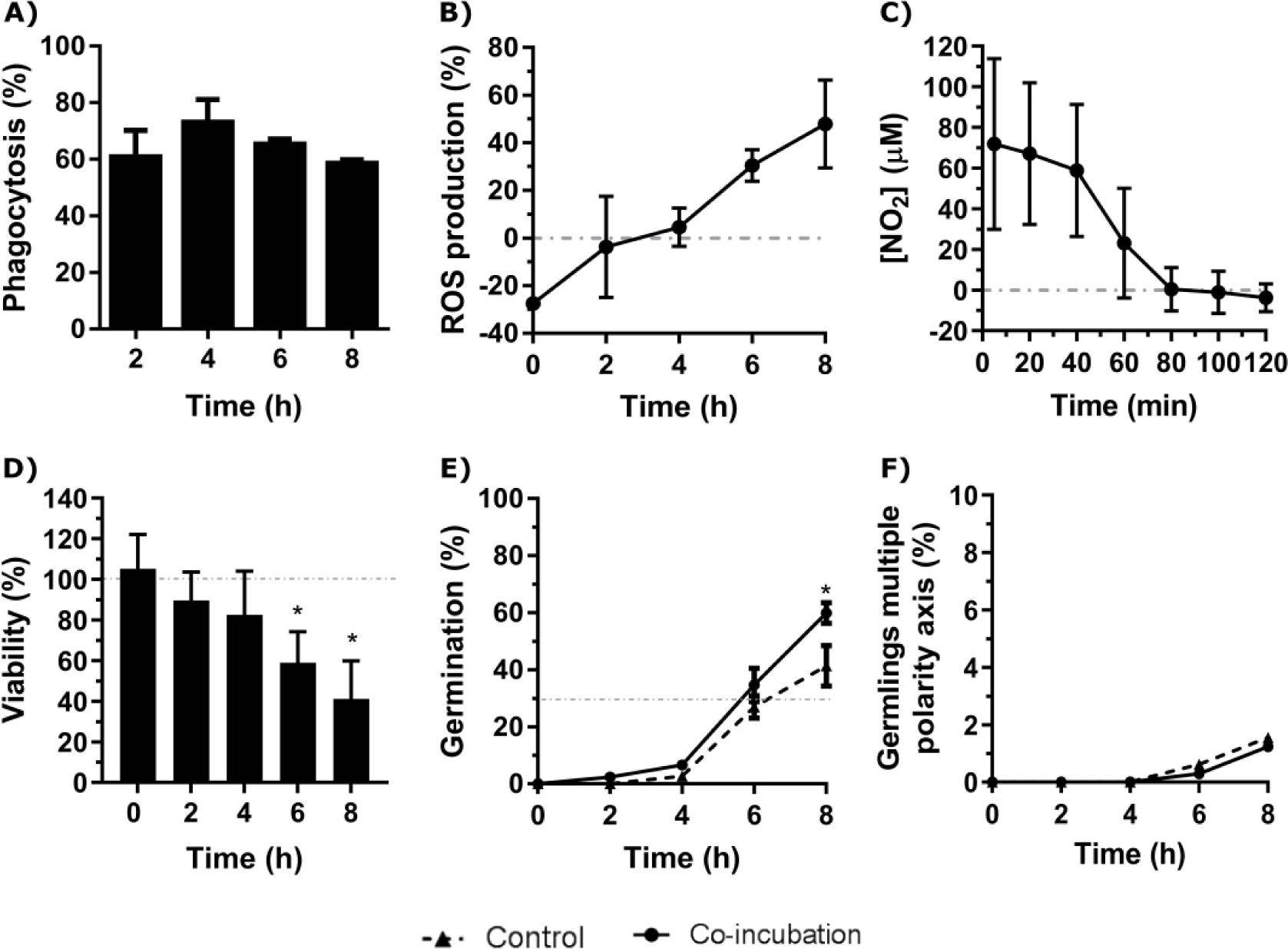
Characterization of the co-incubation between the fungal strain Af293 and the macrophage cell line RAW 264.7. **A)** Phagocytosis assay during 8 hours of co-incubation with Af293. **B)** Reactive oxygen species (ROS) production of the RAW 264.7 macrophages during co-incubation with Af293. The dashed line corresponds to ROS production by control macrophages. **C)** NO_2_ production of RAW 264.7 macrophages in co-incubation with Af293. The dashed line corresponds to RNS production by control macrophages. **D)** Viability of Af293 after incubation with RAW 264.7 macrophages. The results are relative to the Af293 strain growing without macrophages. The dashed box indicates 100% of cellular viability. **E)** Percentage of germination of the Af293 strain alone and in with co-incubation with RAW 264.7 macrophages. The dashed line indicates 30% germination. **F)** Percentage of Af293 germlings with multiple polarity axes alone and in co-incubation with RAW 264.7 macrophages. *p < 0.05

### Study of co-incubation between *A. fumigatus* and epithelial cells

Co-incubation of *A. fumigatus* with the lung epithelial cell line, A549, revealed a maximum endocytosis of approximately 5% after 6 hours of co-incubation (Fig. 2A). In addition, the contact with these epithelial cells generated a slight inhibition of the germination rate. Indeed, 30% of germination was reached after 8 hours of co-incubation with the epithelial cells while this germination rate was reached at 6 hours when the fungus grew alone (Fig. 2B). However, an increase in germlings with multiple polarity axis was observed after 8 hours of co-incubation of *A. fumigatus* with epithelial cells (Fig. 2C).

**Figure 2.**
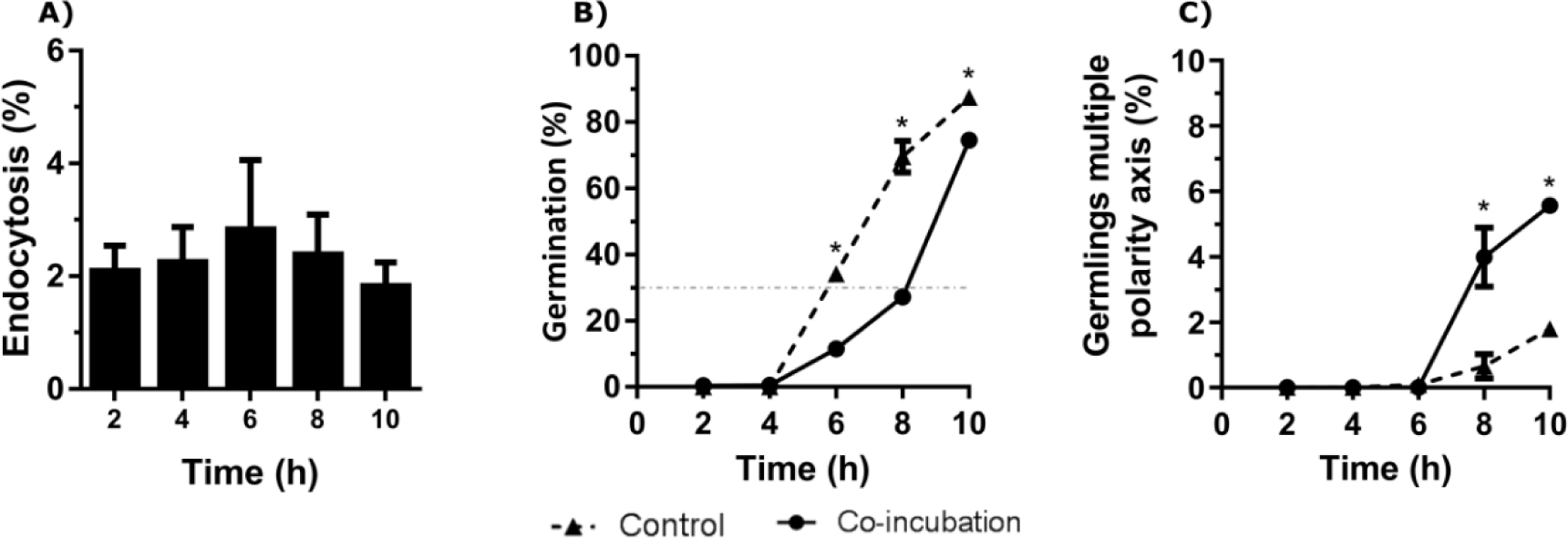
Characterization of the co-incubation between the fungal strain Af293 and the human epithelial cell line A549. **A)** Endocytosis assay during 10 hours of co-incubation with Af293. **B)** Percentage of germination of Af293 in co-incubation with the lung epithelial cell line A549. The dashed line indicates 30% germination. **C)** Percentage of Af293 germlings with multiple polarity axes alone and in co-incubation with the lung epithelial cell line A549. *p < 0.05.

### *A. fumigatus* gene expression in response to the co-incubation with RAW 264.7 macrophage or A549 epithelial cell lines

After the selection of 6 and 8 hours of co-incubation with macrophages and epithelial cells (around 30% of fungal germination), respectively, three samples of mRNA from each condition obtained in independent experiments, as well as their respective controls, were hybridized with the AWAFUGE microarray. Once the data were analyzed and normalized, 2,137 and 5,325 *A. fumigatus* genes were found differentially expressed with respect to their controls (*A. fumigatus* without cells) when the fungus was incubated with macrophages and with human lung epithelial cells, respectively.

To limit and prioritize the genes to be studied, only those displaying > 1.5 or < −1.5-fold change (log_2_) (FC) in their conditions were considered. By this criteria, 235 *A. fumigatus* genes were down-regulated, and 280 genes were up-regulated during the co-incubation with the macrophages (Fig. 3A), whereas 534 were down-regulated and 878 genes were up-regulated during the contact with the epithelial cells (Fig. 3B).

**Figure 3.**
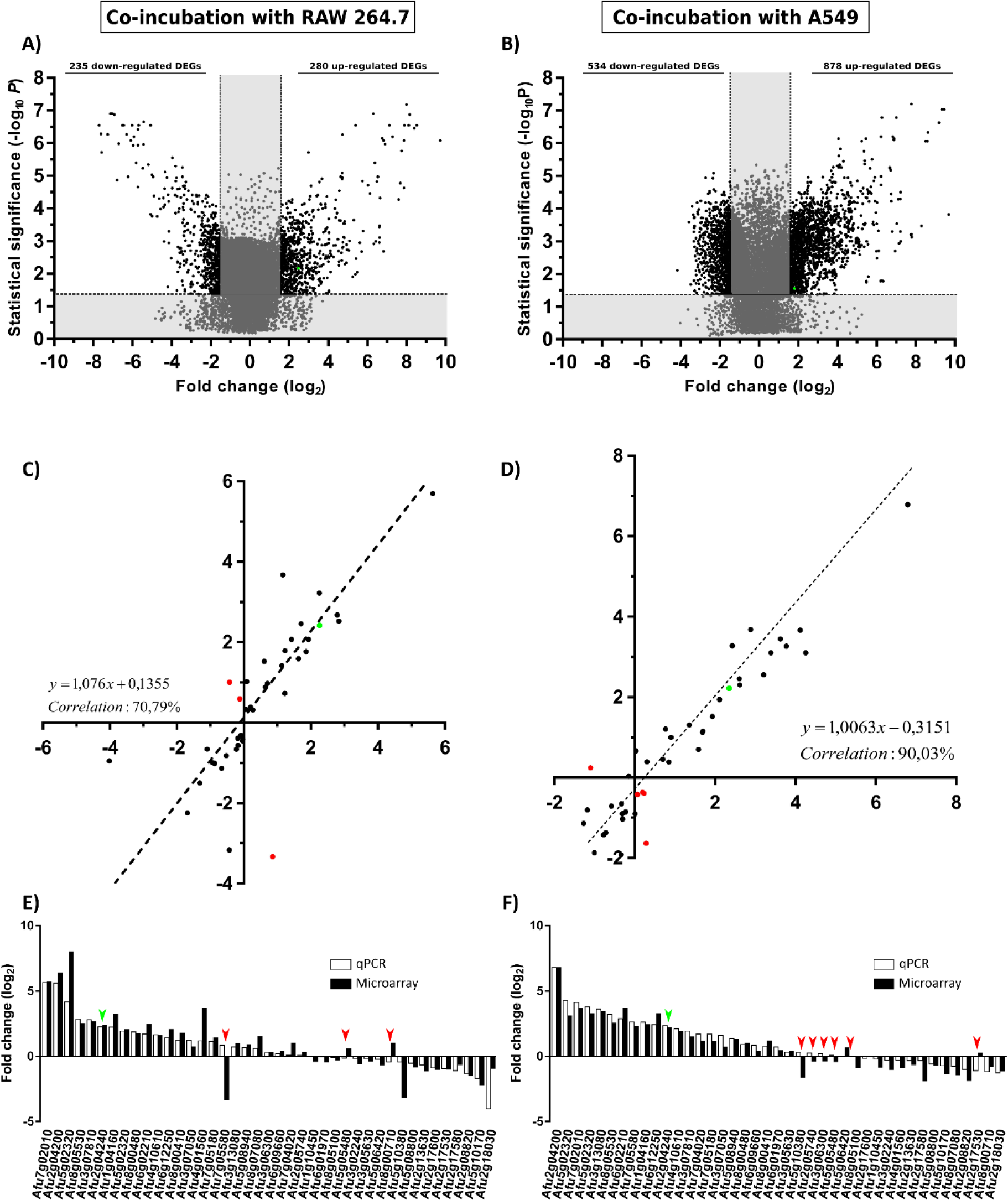
Gene expression analysis. Volcano plot showing *A. fumigatus* differentially (black) (FC > 1.5 or FC < −1.5) and non-differentially (grey) expressed genes in co-incubation with RAW 264.7 macrophages (**A**), A549 lung epithelial cells. (**B**). The x-axis values represent the fold change (log_2_) of microarray data and the y-axis values represent the statistical significance (–log_10_P). Spots with positive values indicate upregulation of the gene during co-incubation with the indicated cell line. **C** and **D)** Correlation analysis between microarray and RT-qPCR data. The x-axis values represent the fold change (log_2_) of microarray data and the y-axis values represent the fold change (log_2_) of RT-qPCR results of the selected fungal genes in co-incubation with RAW 264.7 macrophages (**C**), and A549 lung epithelial cells (**D**). Each point corresponds to the mean value from three independent samples. **E)** Comparative levels of fungal gene expression between the microarray and the RT-qPCR (Af293 in co-incubation RAW 264.7 macrophages). **F)** Comparative levels of fungal genes expression between the microarray and the RT-qPCR (Af293 in co-incubation with A549 lung epithelial cells). Green points in **A**, **B**, **C** and **D** and green arrows in **E** and **F** show the corresponding results obtained for *maiA*. Red spots in **C** and **D** and red arrows in **E** and **F** show contradictory results in gene expression detected between microarrays and RT-qPCR results.

The transcriptomic data obtained from the AWAFUGE microarray was confirmed by RT-qPCR, showing a good correlation result between both techniques. Specifically, a correlation of 70.79% and 90.03% was obtained between microarray and RT-qPCR verification in the co-incubation of *A. fumigatus* with macrophages, and with epithelial cells, respectively (Fig. 3C and 3D). In Fig. 3E and 3F, FC values obtained for each gene and technique used in the validation process are shown. All of them, except those marked in red (dots in panels C and D and arrows in panels E and F), displayed a similar expression in both techniques. In addition, *maiA* gene expression values were plotted in green.

### GO enrichment analysis of the most up/down-regulated DEGs and comparison between different infection models

To identify those *A. fumigatus* processes, components, and functions of greatest importance that were impacted during the experimental infection procedure, we performed a Gene Ontology (GO) enrichment analysis (the complete analysis of DEGs can be found in Table S1).

Among the DEGs, 227 (RAW 264.7 vs Control) and 592 (A549 vs Control) were associated with a known biological process. The five most relevant processes in this category are those related with transport, regulation, response to stress, secondary metabolic process, and lipid metabolic process.

In addition, 252 *A. fumigatus* genes (RAW 264.7 vs Control) and 605 genes (A549 vs Control) were associated with a known molecular function. The categories corresponding to the most genes are oxidoreductase activity, hydrolase activity, and transporter and transferase activities.

Moreover, it is remarkable that 321 (RAW 264.7 vs Control) and 795 (A549 vs Control) genes have been previously associated with a known cellular location, including (in order of gene numbers associated) membrane, mitochondrion, nucleus, cytosol, plasma membrane and the extracellular region.

To select the most important genes related to infection, among the large amount of overexpressed *A. fumigatus* genes found, those up-regulated with a FC > 1.5 in co-incubation in common between macrophages and epithelial cells were selected. Using this approach, a total of 140 *A. fumigatus* genes were obtained (Table S2). These genes were then cross-referenced with those detected as overexpressed during a previous mouse infection model (Guruceaga et al., 2018). This analysis decreased the list to 13 significantly up-regulated summarized in Table 1. Among those, Afu2g04240 (*maiA*) gene was selected as a good candidate to be characterized in this study for its role in *A. fumigatus* for the next reasons: i) Is one of the genes related with the Phe/Tyr degradation pathway still unstudied, ii) Its potential role in the infection process detected after the transcriptomic analysis above explained.

**Table 1.**
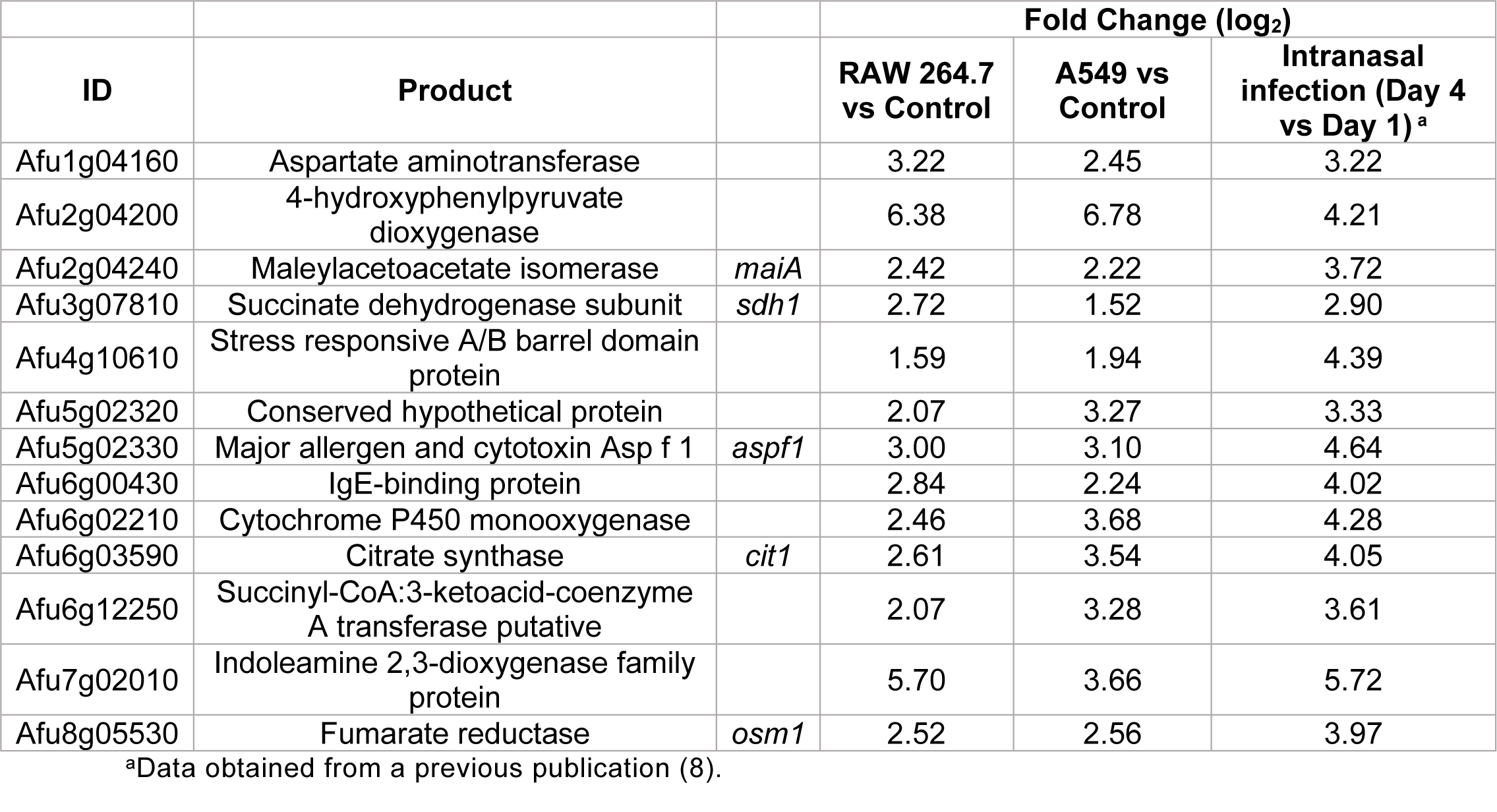
Common overexpressed DEGs in the three experimental infection models.

| ID | Product |  | Fold Change (log<sub>2</sub>) |  |  |
| --- | --- | --- | --- | --- | --- |
|  |  |  | RAW 264.7<br>vs Control | A549 vs<br>Control | Intranasal<br>infection (Day 4<br>vs Day 1)<sup>a</sup> |
| Afu1g04160 | Aspartate aminotransferase |  | 3.22 | 2.45 | 3.22 |
| Afu2g04200 | 4-hydroxyphenylpyruvate<br>dioxygenase |  | 6.38 | 6.78 | 4.21 |
| Afu2g04240 | Maleylacetoacetate isomerase | *maiA* | 2.42 | 2.22 | 3.72 |
| Afu3g07810 | Succinate dehydrogenase subunit | *sdh1* | 2.72 | 1.52 | 2.90 |
| Afu4g10610 | Stress responsive A/B barrel domain<br>protein |  | 1.59 | 1.94 | 4.39 |
| Afu5g02320 | Conserved hypothetical protein |  | 2.07 | 3.27 | 3.33 |
| Afu5g02330 | Major allergen and cytotoxin Asp f 1 | *aspf1* | 3.00 | 3.10 | 4.64 |
| Afu6g00430 | IgE-binding protein |  | 2.84 | 2.24 | 4.02 |
| Afu6g02210 | Cytochrome P450 monooxygenase |  | 2.46 | 3.68 | 4.28 |
| Afu6g03590 | Citrate synthase | *cit1* | 2.61 | 3.54 | 4.05 |
| Afu6g12250 | Succinyl-CoA:3-ketoacid-coenzyme<br>A transferase putative |  | 2.07 | 3.28 | 3.61 |
| Afu7g02010 | Indoleamine 2,3-dioxygenase family<br>protein |  | 5.70 | 3.66 | 5.72 |
| Afu8g05530 | Fumarate reductase | *osm1* | 2.52 | 2.56 | 3.97 |

<sup>a</sup>Data obtained from a previous publication (8).

### The *maiA* gene is essential for Phe/Tyr degradation pathway

The genetic manipulation strategy followed to generate the mutant strains is described in Materials and Methods section and summarized in Fig. 4. Briefly, a disruption mutant was generated by replacing the initial methionine (iMet) with the hygromycin resistance gene. The resulting mutant strain (*maiA-1*) was resistant to hygromycin and has the target gene silenced due to the lack of the iMet (Fig 4A). The complete deletion strain (Δ*maiA*) was also performed, through complete substitution of the *maiA* locus by the hygromycin resistance gene (Fig 4B). Finally, the complement of the *maiA-1* strain (*maiA-1^comp.^*) was constructed by replacing the disrupted allele with the native *maiA* followed by the phleomycin resistant gene in the native locus (Fig. 4C). All the strains were generated using CRISPR-Cas9 highly efficient genetic technology in the Af293 genetic background. The Δ*maiA* strain was constructed to validate the results obtained in the disruption strain. As is possible to see in (Fig. S1 and S2) the deletion strain phenocopied the *maiA-1* strain results, confirming the usefulness of the disruption strategy.

**Figure 4.**
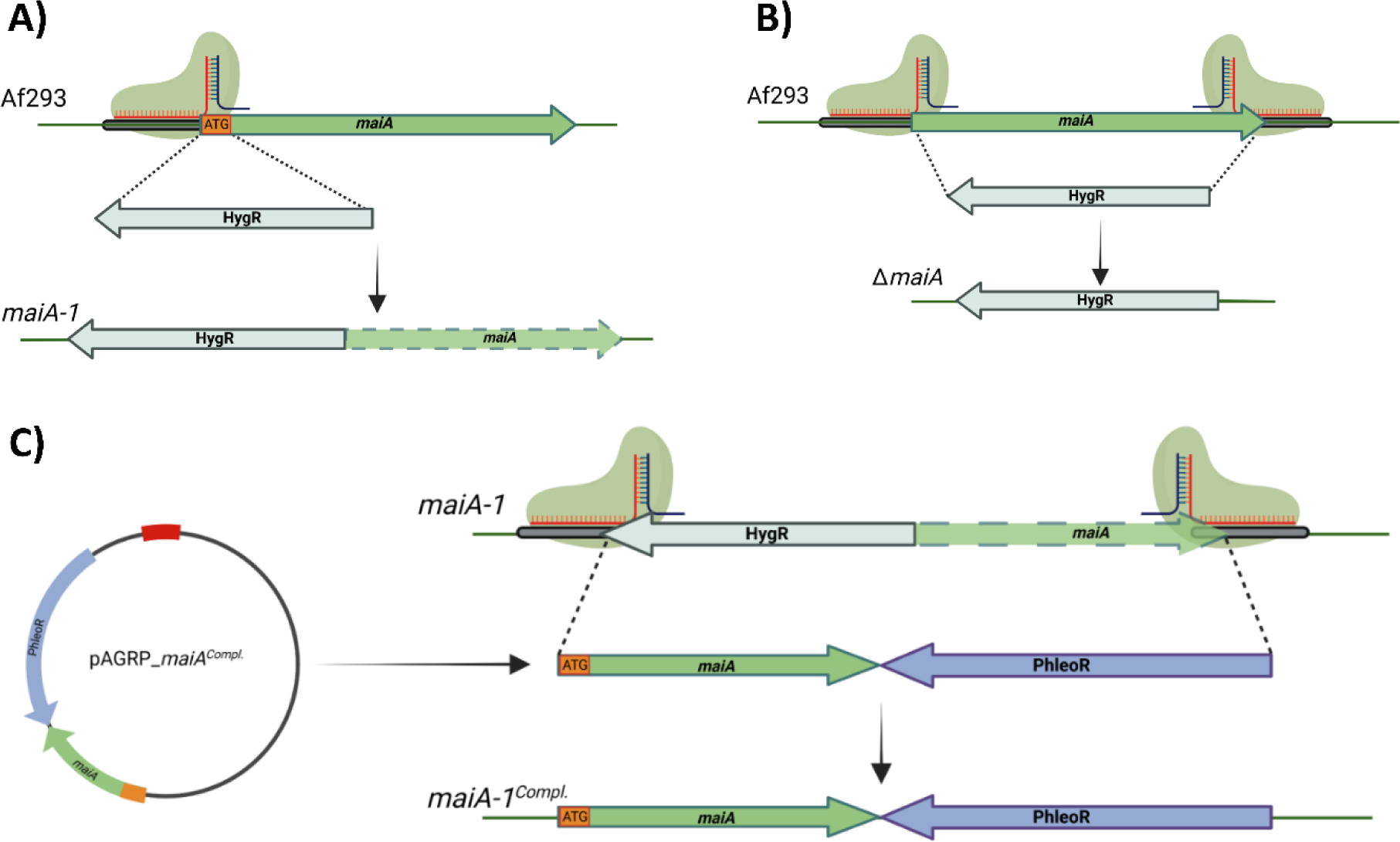
Schematic of gene manipulations by CRISPR/Cas9 editing. **A)** Silencing strategy of the *maiA* gene. One protospacer adjacent motif was designed to disrupt the iMet (ATG orange square) of the *maiA* gene. The disruption was repaired using the hygromycin resistance cassette with 40 base pair microhomology regions upstream and downstream the *maiA* gene in the Af293 genetic background to generate the *maiA-1* mutant strain (*maiA* green dashed arrow represents non-functional *maiA* gene). **B)** The complete deletion of the *maiA* gene was carried using two protospacers adjacent motif designed to the flanking regions of the *maiA* gene. The repair template process was developed following the same strategy described in panel **A** to generate the Δ*maiA* mutant strain. **C)** To complement the *maiA-1* mutant strain, the native *maiA* gene was cloned into pAGRP plasmid (iMet included) upstream the phleomycin resistant cassette and generating the pAGRP_*maiA^comp.^* plasmid. Two protospacers adjacent motif designed to the flanking regions of the *maiA* gene were used. The repair template cassette was amplified from the pAGRP_*maiA^comp.^* plasmid using primers containing 40 base pair microhomology regions upstream and downstream the HygR-*maiA* disrupted regions, generating the *maiA^comp.^* mutant strain. All the colonies were confirmed by PCR and sequencing. The genetic maps were created with BioRender.com.

As mentioned above, we hypothesized that loss-of-function mutations at the *maiA* locus would produce high concentrations of 4-MA that could polymerize spontaneously to pyomelanin (Fig. 5A). To study the pyomelanin production ability, Af293 as well as the *maiA-1* mutant and the *maiA-1^comp.^* strains were grown for 72 hours in GMM broth (Fig. 5B), GMM broth supplemented with 10 mM of Phe (Fig. 5C) or GMM broth supplemented with 10 mM of Tyr (Fig. 5D). Indeed, when Phe or Tyr were added to the media, the *maiA-1* mutant strain accumulated pyomelanin indicated by an increase of the 405 nm absorbance signal (Fig. 5C) that can be also visually observed (Fig. 5D). A varying ability to degrade both amino acids was shown since the production of pyomelanin was different when Phe or Tyr were used, likely because Phe may be utilized by other metabolic pathways compared to Tyr (Fig. S3).

**Figure 5.**
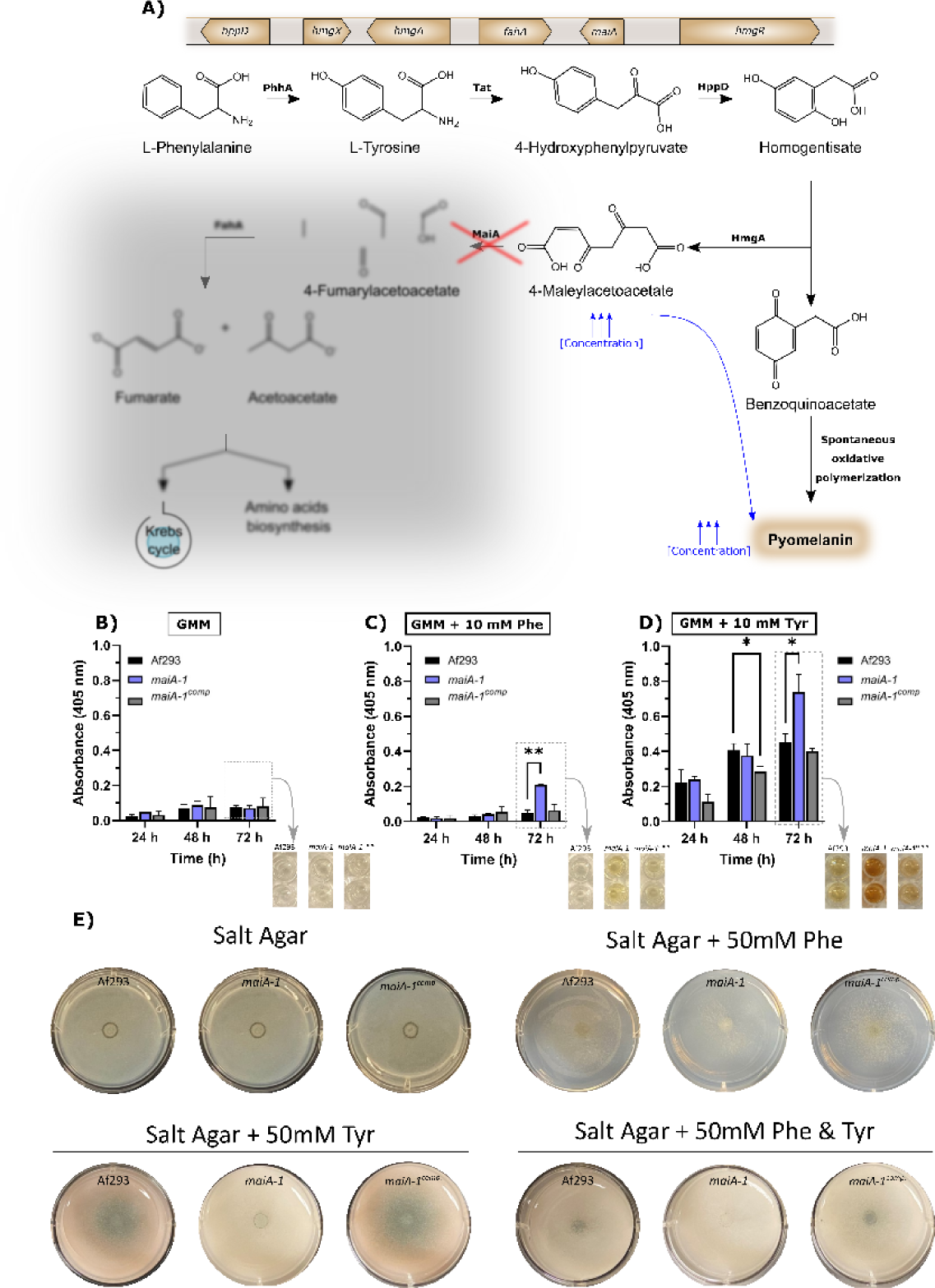
Fungal strain characterization results in relation to pyomelanin production and growth ability in presence of Phe and/or Tyr. **A)** Phe degradation pathway including the genetic cluster and the proteins involved in each step. The red cross indicates the point in which the pathway is disrupted due to the knock down mutation produced in *maiA-1* strain. The diffused grey area indicates the part of the pathway that the *maiA-1* strain cannot perform. Pyomelanin secretion ability of Af293 and *maiA-1* growing in **B)** GMM broth, **C)** GMM broth supplemented with 10 mM Phe and **D)** GMM broth supplemented with 10 mM Tyr. Dashed boxes in **B**, **C** and **D** indicate quantitative information corresponding to the photographs indicated by arrows. **E)** Spotting assay after 72 hours of incubation of the Af293 and *maiA-1* strains on salt agar plates in which the only carbon source was Phe (50 mM), Tyr (50 mM), or Phe and Tyr (50 mM each).

It was also striking that when salt agar plates were supplemented with 50 mM of Phe, 50 mM of Tyr or 50 mM of both amino acids as the sole carbon source, the *maiA-1* strain exhibited impaired growth (Fig. 5E). Attempts to measure biomass accumulation in submerged culture to further quantify the reduced ability to utilize these amino acids as carbon sources revealed a complete lack of growth for the *maiA-1* mutant in both Phe and Tyr (data not shown).

### Role of *maiA* in growth and development of *A. fumigatus*

To understand the role and importance of the *maiA* gene in the basal growth of *A. fumigatus*, the *maiA-1* disrupted strain was analyzed using germination and radial growth assays in GMM). Although the germination rate of all the strains was the same over the time studied (Fig. 6A), slight differences in the radial growth ability were found between the Af293, the *maiA-1* and *maiA-1^comp.^* after 72 hours of growth. After 96 hours, no difference in the radial growth between Af293 and the complemented strain was found. In contrast the *maiA-1* mutant strain colonies were slightly smaller compared to those from the Af293 background (Fig. 6B). Although statistical differences exist at the various timepoints, no overt defects in colony formation were seen.

**Figure 6.**
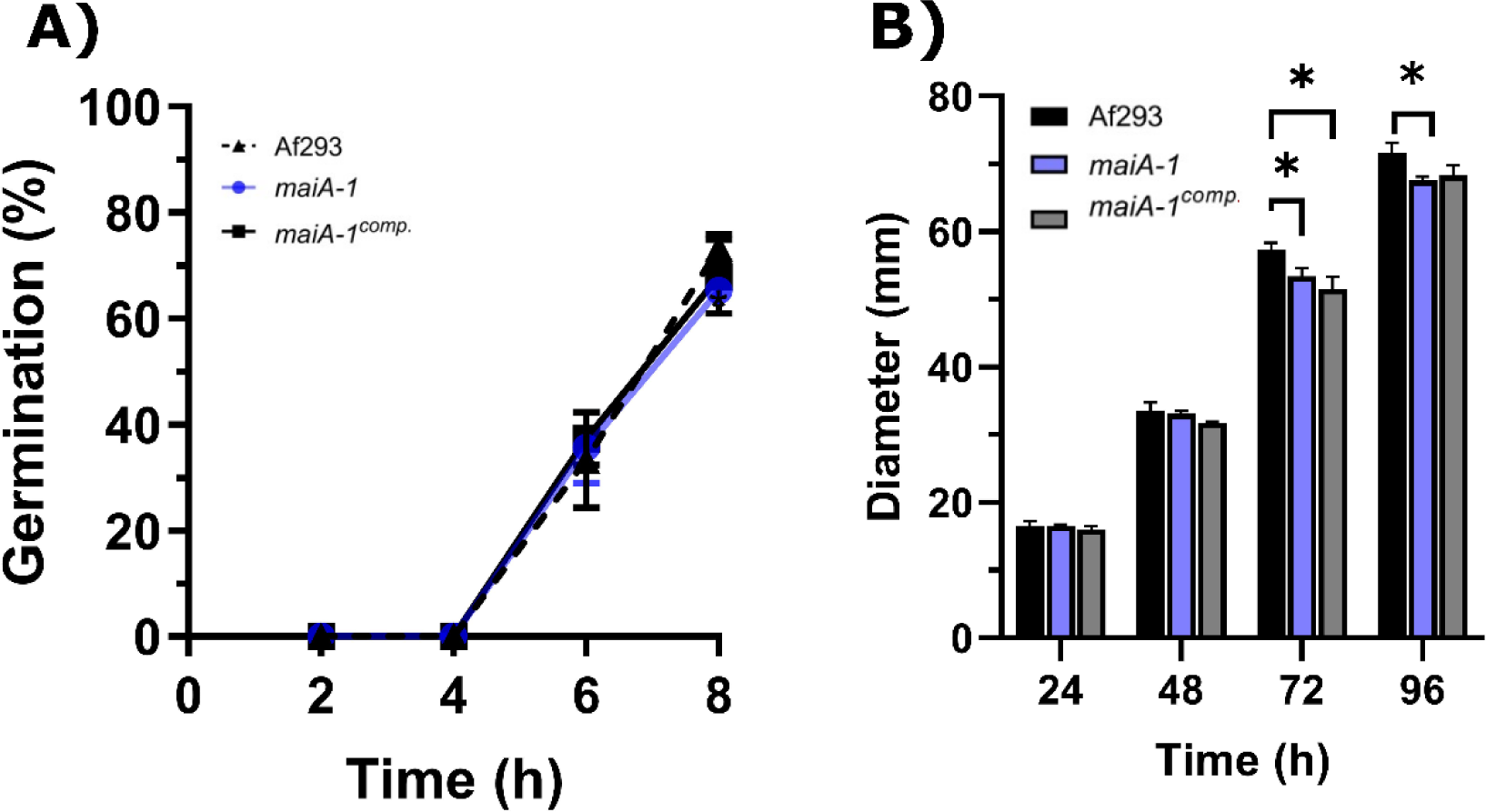
Cellular characterization results of the *maiA-1 mutant* strain. **A)** Percentage of germination during 8 hours of incubation in GMM of the Af293, *maiA-1*, and *maiA-1^comp.^* strains. **B)** Diameter evolution of the colonies of both strains in GMM plates. *p< 0.05.

To better understand the role of *maiA* in maintaining fungal cell wall homeostasis, a stress response assay using two cell wall stressor compounds was performed (Fig. 7). The *maiA-1* mutant strain was hyper-susceptible to the cell wall stressors congo red (CR) and calcofluor white (CFW) as observed by the inability to grow at concentrations higher than 40 µg/ml of both compounds in GMM (Fig. 7 top panels). The ability of this mutant strain to grow in the presence of both compounds was recovered by the addition of sorbitol to the media (SMM) (Fig. 7 bottom panels), further supporting that growth inhibition in the mutant was due to cell wall instability.

**Figure 7.**
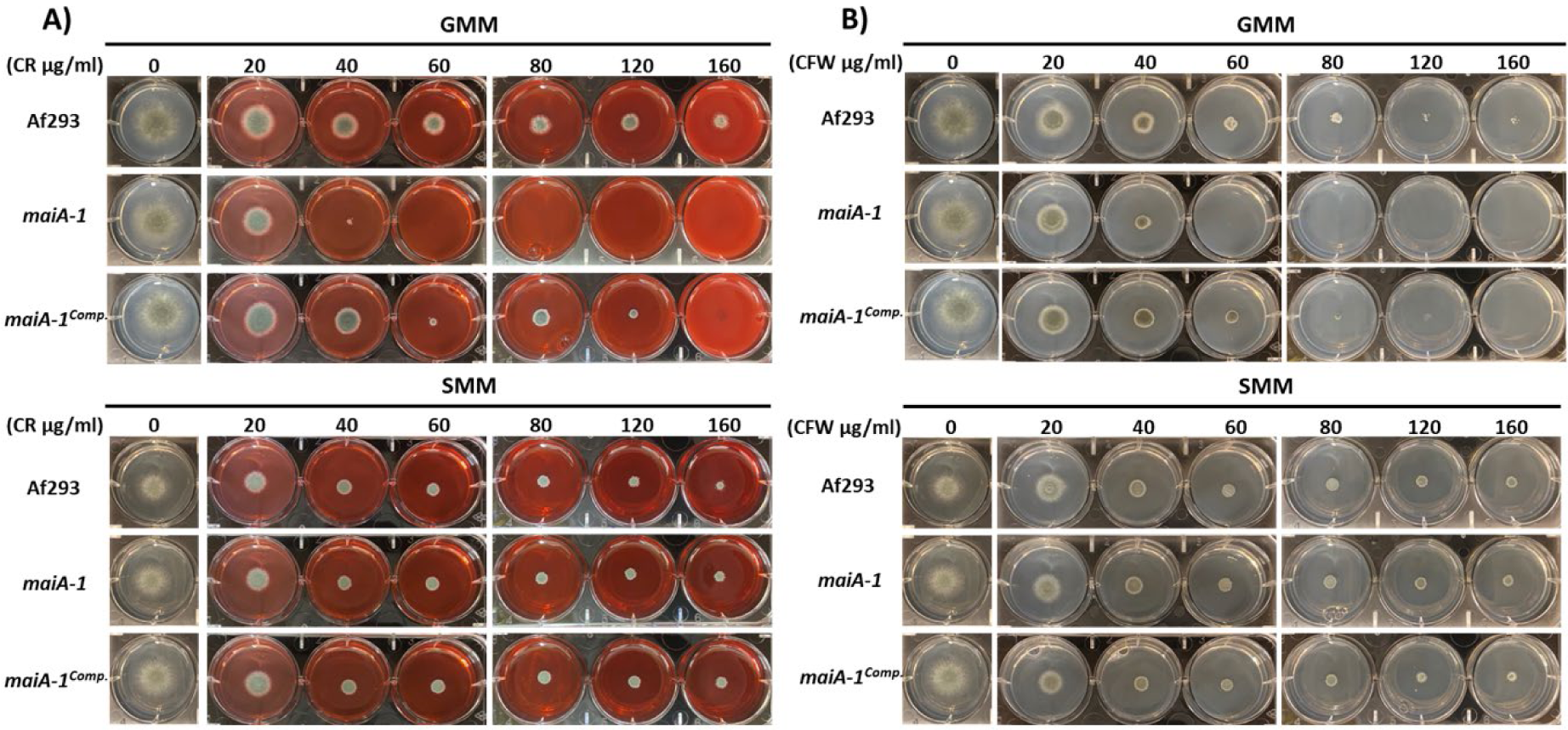
Phenotypic characterization of Af293, *maiA-1*, and *maiA-1^comp.^* strains using GMM agar 6-well plates. GMM or SMM 6 well plates supplemented with different concentrations (0, 20, 40, 60, 80, 120, 160 µg/ml) of congo red (CR) (**A**) or calcofluor white (CFW) (**B**).

### The *maiA-1* hyphae display an unstructured morphology

Scanning electron microscopy (SEM) analysis of the mycelial appearance after culturing for 12 hours in GMM broth showed that the Af293 strain maintained a normal hyphal morphology with typical 45° branch angles and mycelia that were strongly attached to the surface (Fig. 8A). In addition, analyzing the images at higher magnification, the Af293 hyphae were surrounded by an extracellular matrix, which was more prevalent at the hyphal apex (Fig. 8C) On the contrary, the *maiA-1* strain displayed a disorganized mycelium, which seemed to be more aerial, less attached to the surface and in consequence more rounded (Fig. 8B) without the surrounding matrix observed in Af293 (Fig. 8D).

**Figure 8.**
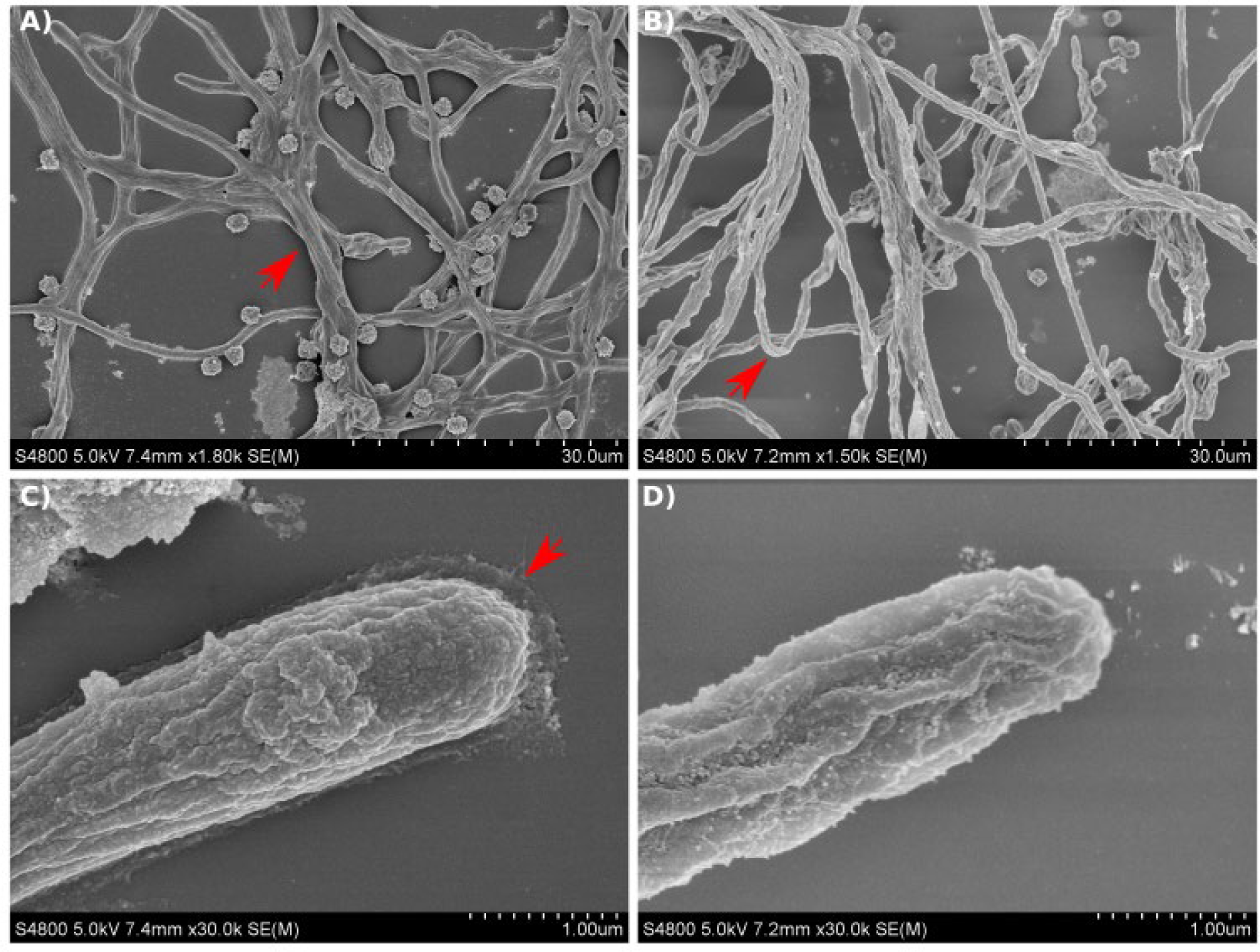
Scanning electron micrographs of *A. fumigatus* strains growing in GMM broth for 12 hours. **A and C)** SEM images of Af293 hyphae. **B and D)** SEM images of *maiA-1* hyphae. Red arrows denote specific details described in text.

### *maiA* is required for normal *A. fumigatus* virulence

Finally, we characterized the pathogenic abilities of the *maiA-1* strain with a set of *in vitro* and *in vivo* assays. First, we performed an *in vitro* characterization challenging mouse bone marrow-derived macrophages (BMMs) with our isogenic set of strains. No differences in phagocytosis rates were observed over 8 hours of co-culture between the BMMs and any of the three fungal strains (Fig. 9A). These results could be related with the H_2_O_2_ susceptibility assay in which similar sized zones of clearance were found in all strains (Fig. 9B), indicating a likely identical resistance to killing inside the phagolysosome. In a detailed study of the BMM cells-pathogen interaction, significantly less TNF release was observed in the macrophages co-cultured with the *maiA-1* mutant strain compared to those co-cultured with either the wild-type or *maiA-1^comp.^* strain (Fig. 9C). A very interesting finding was discovered when the β-glucan content of each fungal strain was studied. Specifically, the total glucan content was found to be significantly lower in the *maiA-1* mutant strain compared to the wild-type strain (Fig. 9D). This result might indicate that reduced stimulation of BMMs induced by the *maiA-1* mutant strain could be attributable to its lower β-glucan content.

**Figure 9.**
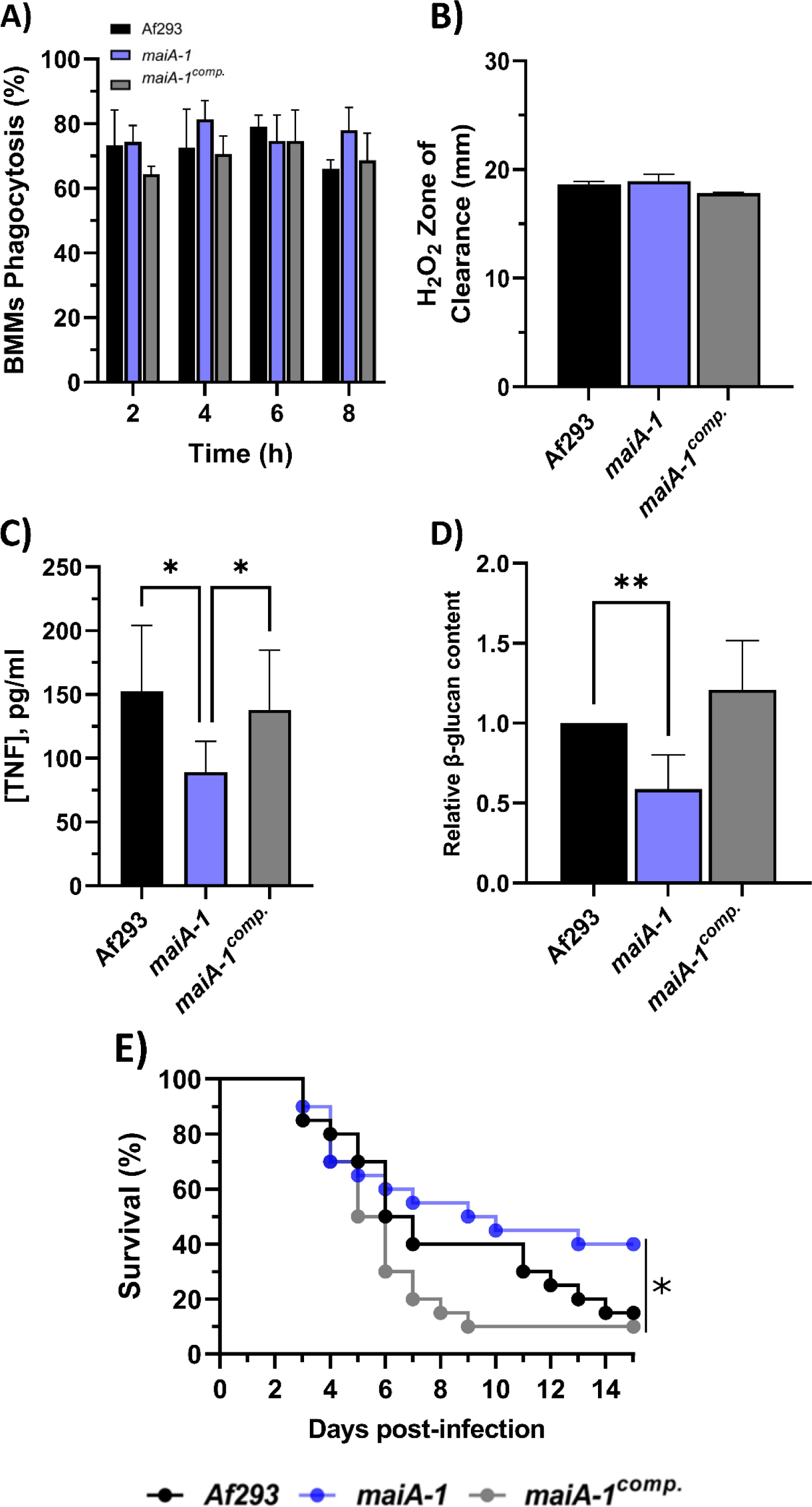
Characterization of the Af293, *maiA-1*, and *maiA-1^comp.^* strains during the infection process. **A)** Phagocytosis measured after 8 hours of co-incubation between the three fungal strains and primary BMMs. **B)** Inhibition halus diameter of a culture of the three fungal strains on GMM agar plates after 48 hours of incubation in presence of 200 mM H_2_O_2_ in a central well of the plate. **C)** BMMs TNF release after 16 hours challenge with the three fungal strains. **D)** Relative content of β-glucans quantified in the three fungal strains. **E)** Kaplan-Meier analysis of chemotherapeutically immune suppressed infected mice for 15 days post intranasal exposition to Af293, *maiA-1*, and *maiA-1^comp.^* fungal strains. *p< 0.05.

Moreover, the impact of the absence of the *maiA* gene on virulence was studied using a neutropenic mouse model of invasive aspergillosis (IA) (Fig. 9E). The results revealed that 85% and 90% of the mice infected with the Af293 strain and the *maiA-1^comp.^,* respectively, had died after 15 days post-infection. In contrast, the group of mice infected with the *maiA-1* strain showed 40% survival at the end of the assay. The overall statistical analysis of the assay showed a significantly lower lethality of the *maiA-1* mutant strain.

## Discussion

In this work, RAW 264.7 macrophages and A549 human lung epithelial cells were used as *in vitro* infection models in co-incubation with *A. fumigatus* to perform two independent transcriptomic studies. After characterization of both models during co-incubation, when *A. fumigatus* reached 30% germination, the total mRNA was isolated and hybridize with the AWAFUGE 1.0 microarray. The transcriptomic analysis of both *in vitro* experimental assays showed that 140 *A. fumigatus* were up-regulated (FC > 1.5) in common in the two models used. To further prioritize the number of genes of interest, they were then compared with those detected as up-regulated during a previously published murine intranasal infection (8). Using this approach, 13 genes were found to be up-regulated in all three abovementioned infection procedures. Some of these genes have been previously studied by other authors and have been related with the infection process or detected as expressed at the onset of the contact between cells or sera. Among them, it is important to highlight genes such as *aspf1*, that codifies an *A. fumigatus* allergen commonly found in different proteomic/transcriptomic studies (8, 9) or the fumarate reductase Osm1, whose expression has been previously observed during hypoxia and during the first contact between the dormant conidia and the host (10, 11). Two up-regulated genes codify TCA cycle enzymes: the succinate dehydrogenase Sdh1 and the citrate synthase Cit1. The former protein has been previously described as expressed after human neutrophil exposure (12), during the exit of the dormancy (13) as well as under hypoxia (11). Regarding the Cit1, some researchers have detect its expression during the contact with sera (14) and with airway epithelial cells (15); in addition, the *cit1* gene was also described as essential for IA manifestation (16).

The *A. fumigatus* catabolic Phe/Tyr degradation pathway is encoded by a cluster of 6 genes (*hppD*, *hmgX*, *hmgA*, *fahA*, *maiA*, and *hmgR*) in this microorganism (3, 4). Except for two genes (*fahA* and *maiA*), the cluster of genes involved in this pathway has been previously studied through the development of loss-of-function mutant strains. The Δ*hppD* null mutant displayed a lack of pyomelanin synthesis because HGA formation was blocked. Furthermore, the hyphae of this mutant showed increased susceptibility to hydrogen peroxide and other oxidizing agents when compared with Af293 (4). Moreover, this gene has been detected as overexpressed at 96 hours post-intranasal infection in comparison with the first 24 hours in immunosuppressed mice (8). Other authors also detected the overexpression of *hpdA*, the *hppD* homolog in *Penicillium marneffei,* after exposure to macrophages (17). In addition, the Δ*hmgX* mutant strain has been shown to have similarities with a Δ*hppD* strain (18). This Δ*hmgX* mutant was unable to produce HGA and, therefore, HmgX could be a cofactor or mediator of HppD enzyme function (18). On the other hand, the Δ*hmgA* mutant strain showed an increase in pyomelanin owing to the accumulation of HGA and displays reduced susceptibility to oxidizing agents compared to Δ*hppD* (4). Finally, the last gene of the Phe/Tyr degradation pathway studied was the transcription factor *hmgR*. The Δ*hmgR* mutant is incapable of using Tyr as the sole carbon source or nitrogen source and, therefore, only a residual accumulation of pyomelanin was observed in cultures (18).

The only published study concerning *maiA* in *A. fumigatus* showed that this gene was overexpressed when the medium was supplemented with Tyr (18). Orthologs of this gene has been silenced in both *Aspergillus nidulans* and *P. marneffei* (17, 19, 20). The catabolic Phe/Tyr degradation pathway, in which *maiA* is putatively involved, is responsible for the production of one type of soluble melanin, known as pyomelanin, when an accumulation of intermediate metabolites, such as HGA or 4-MA, occurs. The mutation of the *maiA* gene is predicted to result in truncation of the final part of the pathway inhibiting the degradation of Phe or Tyr to fumarate and acetoacetate, which are destined to the Krebs cycle or the biosynthesis of other amino acids. Consequently, there would be an accumulation of 4-MA and HGA and the subsequent production of pyomelanin.

In accordance to previous publications (3, 4, 18), in response to Phe and Tyr, the *maiA-1* mutant strain would block the Phe/Tyr degradation pathway causing a probable accumulation of 4-MA and, likely, of the previous metabolites of the pathway that can polymerizate to pyomelanin. Specifically, the mutant strain *maiA-1* started to produce this pigment when Phe and, especially, Tyr were added to the medium as the only carbon source. These results support our hypothesis that *A. fumigatus* can uses in a more effective way Phe than Tyr as an alternative carbo source. However, results obtained by other investigators using the Δ*hmgA* mutant showed that this strain produced more pyomelanin by the addition of Phe than Tyr (4).

Regarding basal growth analysis, our results suggest that the absence of the *maiA* gene did not impact on germination and minimally impacted on basic radial growth of the colonies. In contrast, the cell wall stress assays carried out in this study pointed out that the *maiA-1* strain suffers an alteration of the cell wall stability, which makes it unable to grow in presence of CR or CW, perhaps due to the lower content of glucans detected in its wall. CR and CW are compounds with a chemical structure that can interact with β-linked glucans and, specifically, it is thought that both compounds interfere with cell wall assembly by binding to chitin. Conceivably, CW and CR act by binding to nascent chitin chains, thereby inhibiting the assembly enzymes that connect chitin to β-1,3-glucan and β-1,6-glucan (21–23). The ability to grow in the presence of both compounds was recovered by osmotic stabilization using sorbitol.

Other studies performed with mutant strains related with high-osmolarity glycerol response pathway (*shoA*, *msbA*, *opyA*, *sakA* and *mpkC*) which reported similar cell wall phenotypes (i.e., more sensitive to cell wall stress) are also more sensitive to osmotic stresses (24, 25) and to caspofungin. In contrast, the *maiA-1* strain displayed the same growth ability as the Af293 strain in the presence of osmotic stresses such as KCl or NaCl. In addition, the *maiA-1* strain showed normal susceptibility to the antifungal drugs tested (voriconazole, fluconazole, posaconazole, caspofungin and micafungin) following the EUCAST standardized method (data not shown). It is true that, while the *maiA* gene is involved in Phe/tyr degradation pathway, the other aforementioned genes are involved in the high-osmolarity glycerol response pathway and, therefore, the studies are not totally comparable. Another possible explanation is that, although the *maiA-1* mutant strain displayed a cell wall defect, this is not caused through an issue in the cell wall integrity pathway because this mutant strain was not affected by the presence of caffeine in the media as other investigators have previously described (5, 26). This issue could be generated by a blockage of the Phe/Tyr degradation pathway since the *maiA-1* strain is unable to complete the catabolic process of Phe and Tyr by which the fungus could synthesize new amino acids for structural purposes.

The SEM analysis showed a general unstructured appearance of the *maiA-1* mutant strain confirming a cell wall issue as we hypothesized from the CR and CFW stress assays. The Af293 hyphae looked similar to other SEM images presented by other investigators previously (27–29). However, the *maiA-1* hyphae showed differences in the external structure of the cell wall and a lack of a putative matrix-like substance that surrounded the apex of Af293 hyphae.

The last step of the research was to study the role of the *maiA* gene in fungal virulence. The cell wall is the main fungal target recognized by the immune cells and, for that, an *in vitro* model of infection using primary BMMs was performed. The *maiA-1* and Af293 strains showed the same H_2_O_2_ resistance ability, similar to the results previously described using Δ*hppD* and Δ*hmgA* mutants (4). These results could be related to the ability of the fungal cells to survive phagolysosome acidification. Boyce and coworkers performed the same experiment with *P. marneffei* and also found that Δ*hppA* and Δ*hmgA* showed a very mild sensitivity to H_2_O_2_ (17). No other studies have previously described a similar phenotype with alterations in cell wall, and β-glucans composition, upon silencing genes involved in the Phe/Tyr degradation pathway in *A. fumigatus*, *A. nidulans* or *P. marneffei*. However, this result could explain, not only why the *maiA-1* strain produced significantly less TNF release by BMMs after 16 hours, but also the phenotype showed in response to CR and CFW and the SEM images.

The lower mortality observed in neutropenic mice infected with the mutant strain is novel and interesting because other mutant strain of a gene involve in the Phe/Tyr degradation pathway (Δ*hppD*) did not show any difference in the mortality rate observed when was compared with its complemented strain (*hppDc*) (18). In addition, it is important to note that these other studies used only transient immune suppression with corticosteroids instead of our model (cyclophosphamide and triamcinolone acetonide) that employs persistently immunosuppressed mice. The *maiA-1* cell wall defect as well as the reduced pro-inflammatory activity could be the keys to explaining the activity of macrophages against *maiA-1*.

In this study, an involvement of *maiA* with fungal cell wall, which is seemingly independent of classical MAPK regulation, has been established. Furthermore, a new Phe/Tyr fitness theory by which the use of Phe as sole carbon source is energetically more beneficial to *A. fumigatus* is proposed. Finally, the role of the *maiA* gene in fungal virulence has been revealed using a chemotherapeutically immune suppressed murine infection model. In fact, this is the first study in which a mutation of one of the genes involved in the Phe/Tyr degradation pathway is directly related to *A. fumigatus* virulence. The process by which the *maiA-1* strain suffers cell wall defects should be studied in future investigations, but the impossibility of this mutant strain to biosynthesize some amino acids could be involved in the phenotype observed both related with cell wall defects as well as the decreased virulence.

## Material & Methods

### *Aspergillus fumigatus* strains, media, and growth conditions

In this study, we used the *A. fumigatus* Af293 strain as the wild type genetic background to develop the disruption mutant strain *maiA-1* as well as its complement strain *maiA-1^comp.^* and the deletion mutant strain Δ*maiA*. All fungal strains were cultured in Glucose Minimal Media (GMM) at 37°C for five days, and their conidia were harvested directly from the agar plates using sterile water, followed by two washes with saline solution (0.9% NaCl). The concentration of conidia needed for each experiment was calculated using a Bürker counting chamber.

To study the ability of the fungal strains to grow with a sole carbon source, GMM was replaced by a GMM medium without glucose (named salt agar).

### Cell lines

The murine macrophage cell line RAW 264.7 and the human lung epithelium cell line A549, both obtained from the American Type Culture Collection (ATCC, Manassas, VA, USA), have been used in this study. The culture conditions as well as viability calculation and passage method were done following a previously described method (30).

### Phagocytosis assays and fungal behavior against RAW 264.7 cell line

To understand the behavior of the fungal strains in contact with the murine macrophages cell line RAW 264.7 we followed the method previously described (30). Briefly, 2 × 10^5^ cells/ml in 500 µL of cell culture RPMI (RPMI 1640 medium supplemented with 10% heat-inactivated FBS, 200 mM L-glutamine, 100 U/ml penicillin and 0.1 mg/ml streptomycin) were seeded in 24-well plates which contained 12 mm-diameter coverslips. After overnight incubation, RAW264.7 cells were co-cultured with *A. fumigatus* conidia at a multiplicity of infection (MOI) of 10 (ten conidia per cell). In parallel, we seeded the same amount of conidia in the absence of RAW 264.7 cells. After 2-, 4-, 6-, and 8-hours post-incubation we moved the coverslips to a new plate with cold PBS to calculate the percentage of phagocytosis, fungal germination, and hyphal branching using a Nikon Eclipse TE200-U inverted microscope.

### Endocytosis assays and fungal behavior against A549 cell line

To study fungal behavior in contact with the human alveolar epithelial cell line A549 we seeded 1 × 10^6^ cells/ml in 500 µL of RPMI using 24-well plates which contained 12 mm-diameter cover slips. After an overnight incubation, the cells were co-cultured with *A. fumigatus* conidia pre-stained with FITC (conidia were stained overnight at 4°C in an orbital shaker) at a MOI of 5. At each incubation time (2, 4, 6, 8 and 10 hours), we moved the coverslip to a new plate to calculate the percentage of endocytosis. For that, a minimum of 500 cells/conidia/hyphae were counted in each replicate of the experiment using the inverted microscope Nikon Eclipse TE2000-U to calculate endocytosis (%), fungal germination (%) and hyphal branching (%) both in contact with epithelial cell lines and *A. fumigatus* growing alone as the control condition.

### RAW 264.7 ability to produce Reactive Oxygen Species (ROS) and Reactive Nitrogen Species (RNS) against *A. fumigatus*

The ability of RAW 264.7 cells to produce ROS and RNS in response to the fungus *A. fumigatus* was studied in 24-well plates using the same conditions and times described for the phagocytosis assay method.

ROS detection was done using a specific detection kit (Life Technologies, Carlsbad, CA, USA) following the manufacturer’s instructions. Briefly, during co-incubation, the ROS detection reagent was reconstituted by adding 173 μL of DMSO. After that, the vial was diluted 1:100 with sterile PBS and pre-heated in a water bath at 37°C until its use. At the end of the 8-hour infection period, the media was removed from all wells, and we performed 2 washes with sterile, pre-heated PBS. Finally, 500 μL of the pre-heated probe was added. Subsequently, samples were incubated at 37°C, 5% CO_2_ for 20 minutes to allow incorporation of the probe into the cells. Then, the solution containing free probe was removed and replaced by 500 μL of sterile preheated PBS incubating again for another 20 minutes under the same conditions to allow the cells to process and incorporate the probe. Finally, probe fluorescence was measured in a plate reader (Biotek, Synergy HT) at 492 nm.

The RNS production was performed by measuring the nitrite accumulated in the medium. Once the incubation time finished, 150 µL of the supernatant were mixed with 130 µL of distilled water and 20 µL of an SFA-NED solution (1:1) in a 96-well plate. After that, the plate was incubated at room temperature for 30 minutes in dark. Finally, the absorbance was measured in a plate reader (Biotek, Synergy HT) at 548 nm.

Three independent experiments were performed on different days using the ROS or RNS production of a cell culture that grew without presence of fungus as control.

### RNA isolation and purification

For RNA isolation, we collected the cells and fungus using a cell scraper after the incubation time selected in each case (6.5 hours with RAW 264.7 and 8.5 hours with A549). Then, the samples were centrifuged for 1 minute at 14,000 rpm and the pellet was suspended in 1 ml of pre-chilled DEPC sterile deionized water to lyse the mouse/human cells. After that, the samples were centrifuged again in the same conditions abovementioned, the fungal pellet was suspended in 500 µL of pre-chilled DEPC-treated sterile deionized water and transferred to a 1.5 ml tube containing 200 µL of 0.5 mm glass beads (Sigma-Aldrich, St. Louis, MO). The samples were homogenized using the MillMix 20 beat-beater (Technica, Dulles, VA, USA) at 30 Hz for 2 minutes. Finally, we centrifuged the samples as mentioned above, and the supernatant was recovered and transferred to the extraction columns of the RNeasy Plant Mini Kit (Qiagen). The RNA isolation procedure was finished following the manufactureŕs instructions and the RNA quantity and integrity was verified on a 2100 Bioanalyzer (Agilent Technologies, Santa Clara, CA, USA). For microarray analysis and RT-qPCR confirmation, three independent RNA samples for each time point, each of them obtained from an independent co-incubation with cells assays or controls, were studied.

### Microarray selection, hybridization, and expression data analysis

The fungal transcriptome was studied using the AWAFUGE microarray (Agilent Whole *A. fumigatus* Genome Expression 44K v.1). The microarray hybridization and the analysis of the raw data obtained was processed as previously described (8). After the analysis we determined the genes down- or up-regulated relative to the fungal growth alone.

### Microarray data confirmation by reverse transcription quantitative PCR

We selected a subset of 22 genes to verify fungal expression profiles by RT-qPCR co-cultured with RAW 264.7 macrophages, and 22 genes with A549 pneumocytes. In both cases, we used the same 4 reference genes. Specific *A. fumigatus* primers were designed using Primers Quest Tool (available at “https://eu.idtdna.com/site”) to avoid biased results due to mouse or human RNA remaining in the samples (Table S3). Selection of the most suitable housekeeping genes and analysis of the RT-qPCR results were completed following a previous publication (8).

### Gene ontology (GO) analysis

The GO enrichment of those DEGs (log FC > 1.5 or log FC < −1.5), was done using the Hans Knoell Institute FungiFun website (https://elbe.hki-jena.de/fungifun/) (31).

### Gene target selection criteria

The selection of *maiA* as an important *A. fumigatus* gene involved in fungal virulence was determined after analyze our transcriptomic results and previous transcriptomic studies. We classified the genes following their fold change values (FC). We only focused our attention on those genes highly down- or up-regulated (FC < 1.5 or FC > 1.5 respectively). After that, we compared the common *A. fumigatus* up-regulated DEGs in three different assays: 1) *A. fumigatus* strain Af293 in co-incubation with lung epithelial cell line A549, 2) *A. fumigatus* strain Af293 in co-incubation with macrophages RAW 264.7 and 3) *A. fumigatus* strain Af293 infecting immunosuppressed mice (8).

### Mutant strains generation

The *A. fumigatus* mutant strains used in this study were performed following the CRISPR/Cas9 strategy previously described (32). In this study, we used the Af293 genetic background to use the same strain used in the transcriptomic studies in which we selected *maiA* as interesting avoiding bias interpretations. Briefly, *A. fumigatus maiA* mutant strains were performed using Hygromycin B (Thermo Fisher, Waltham, MA, USA) as a selection marker. For the disruption mutant strain (*maiA-1*), Cas9 targets and the corresponding crRNA were designed to delete the initial methionine of the *maiA* gene. In contrast, for Δ*maiA* we designed two gRNAs located in the 5’UTR and 3’UTR respectively with the aim of replacing the whole target gene with the Hyg^R^ cassette. Transformation of *A. fumigatus* protoplasts was carried out following the classic protocol described in the literature (33). Finally, the complement of the *maiA-1* strain (*maiA-1^comp.^*) was constructed by re-introducing the original *maiA* gene followed by the phleomycin resistance cassette in the native locus.

### Determination of the pyomelanin production

Pyomelanin production was measured following a previously published protocol (4). Briefly, GMM broth (150ml) was inoculated with 1 × 10^7^ conidia of each indicated *A. fumigatus* strain. After 20 h of pre-incubation, Phe or Tyr was added to a final concentration of 10 mM. Aliquots of 500 µL of GMM broth were taken at 24, 48 and 72 hours. Pigment formation was analyzed by direct absorbance measurements at 405 nm of the alkalized supernatants obtained by adding 20 µL of 5 M NaOH per ml of sample and centrifuging them at 16,000 x g for 2 min. Three independent replicates of the experiment were carried out on different days.

### Fungal susceptibility to H_2_O_2_ oxidative agent

The susceptibility of the fungal strains to H_2_O_2_ was measured using GMM agar plates following a diffusion assay. Briefly, conidia of both strains (1 × 10^7^) were seeded and spread evenly over the surface of a GMM agar plate (90 mm petri dishes). Once the fungus was seeded, a central well was generated in the middle of the agar plate using a sterile pipet tip. Finally, 50 µL of a 200 mM solution of H_2_O_2_ was added to the central well. After drying for 5 minutes, the plates were incubated at 37°C for 48 hours. Daily, the diameter of the inhibition halo was measured. Three independent replicates of the experiment were carried out on different days.

### Quantification of cell wall glucan content

The β-glucan assay was performed following methods previously described (34–36). Briefly, 1 × 10^7^ conidia of each strain were grown overnight into 25 ml of GMM broth at 37°C. After 16 hours of growing, hyphae were collected by filtration through Miracloth (Sigma-Aldrich, St. Louis, MO, USA) and washed using 0.1 M NaOH solution. Washed fungal hyphae were lyophilized for 24 hours. Five milligrams of dry hyphae were disrupted in a bead-beater three times (1 minute each) with 1 minute of ice incubation between each. Hyphal powder was resuspended to a final concentration of 20 mg/ml in 1 M NaOH and the solution was incubated at 52°C for 30 min. Fifty microliters of each sample were mixed with 185 µL of aniline blue staining solution (183 mM glycine, 229 mM NaOH, 130 mM HCl, and 618 mg/l aniline blue, pH 9.9) into a 96-well masked fluorescence plate (Invitrogen, Waltham, MA, USA). The sample-containing plates were incubated at 52°C for 30 min, followed by a cool down period of 30 min at room temperature. Fluorescence readings were performed using an excitation/emission wavelength of 405/460 nm respectively. All the experiments were performed in triplicate using three independent *A. fumigatus* cultures, and the results were represented as relative quantification versus the Af293 strain.

### *A. fumigatus* cell wall stress assay, spot dilution assay and radial growth

The cell wall stressor assays were done using 6-well plates (Invitrogen, Waltham, MA, USA). The wells were filled with GMM or SMM supplemented with increasing concentrations (20, 40, 60, 80, 120, and 160 µg/ml) of CR or CFW. In parallel, plates with GMM or SMM without stressors were used as controls. To inoculate the plates, 5 µL of a 2 × 10^6^ conidia/ml stock of each strain was pipetted at the center of the plates. The spot dilution assay of each strain was done following the method previously described (37). Briefly, we inoculated 10^4^, 10^3^, and 10^2^ conidia (5 µL/drop) in GMM plates supplemented with 1.2M NaCl or 1.2M KCl as cell membrane stressor compounds or 5 mM Caffeine as a MAPK-inducing stress agent. To study the ability of the strains to use Phe, Tyr, or Phe/Tyr as the sole carbon source, we supplemented salt agar plates with 50 mM Phe, 50 mM Tyr, or 50 mM Phe and 50 mM Tyr. Furthermore, we characterized the radial growth ability of the *maiA-1* mutant strain compared to the Af293. For that, we seeded a 20 µL drop containing a suspension of 10^8^/ml fresh conidia in the middle of a GMM agar plate. The plates were incubated at 37°C for five days and the radial growth of the macroscopic colonies were measured daily.

Three independent replicates of the experiment were carried out on different days. Representative pictures of each condition are shown.

### Scanning electron microscopy (SEM)

For the cell wall surface study, 5 × 10^5^ conidia/ml from the Af293 and *maiA-1* strains were seeded in 1 ml of GMM in 24-well plates containing glass coverslips. The plates were incubated at 37°C, 5% CO_2_ and 95% of humidity for 12 hours. The fungal surface of each strain was studied after 2, 4, 8 and 12 hours of incubation by SEM. At each timepoint indicated, the GMM was removed, and fixing solution (2% glutaraldehyde in PBS buffer) was added to each well for 1h at 4°C. Then the samples were dehydrated through increasing ethanol concentrations and hexamethyldisilane. Finally, they were covered with gold under an argon atmosphere, and visualized under the scanning electron microscope (Hitachi S-4800).

### Mouse bone marrow macrophages (BMMs) isolation and cytokine measurements

BMMs were isolated following previous publications (38). The *in vitro* challenge of the BMMs was done following the same conditions previously described for RAW 264.7 cells. TNF cytokine determination was doing using the mouse TNF-αuncoated ELISA kit (Invitrogen, Waltham, MA, USA) following manufacturer’s instructions after 16 hours of co-incubation between BMMs and *A. fumigatus* in the same conditions previously described for RAW 264.7 and A549 challenges.

### Murine model of invasive pulmonary aspergillosis

Groups of 20 female CD-1 mice (Charles River, Wilmington, MA, USA), weighing approximately 25 g, were immunosuppressed by intraperitoneal injections of 150 mg/kg cyclophosphamide (Sigma-Aldrich, St. Louis, MO) starting 4 days before the inoculation and every 3 days following, using 75 mg/kg of cyclophosphamide, and a single subcutaneous injection of 40 mg/kg triamcinolone acetonide (Kenalog, Bristol-Myers Squibb) 24 hours before the infection. On day 0, mice were transiently anesthetized with isoflurane and challenged via nasal inoculation with 10^6^ conidia in sterile saline solution. Survival was monitored at least twice a day and those animals showing severe signs of distress were humanely euthanized by anoxia with CO_2_. Survival curves were compared using the log-rank test in GraphPad Prism v. 8.2.1 for Windows. The studies were performed in accordance to approved ethical protocols by the LACU committee of the University of Tennessee Health Science Center (Protocol number 22-0373).

### Statistics

All the assays were done in triplicate on three independent days. All the statistical analysis were carried out using GraphPad Prism v. 8.2.1 (GraphPad Software Inc., San Diego, CA, USA) for Windows. In each assay, at least three biological replicates were done to measure each parameter in each condition, avoiding biased results. ANOVA or t-test was used depending on if we did multiple comparisons or compared punctual data, respectively, after ensuring the data sets followed a normal distribution.

---

**Figure S1.**
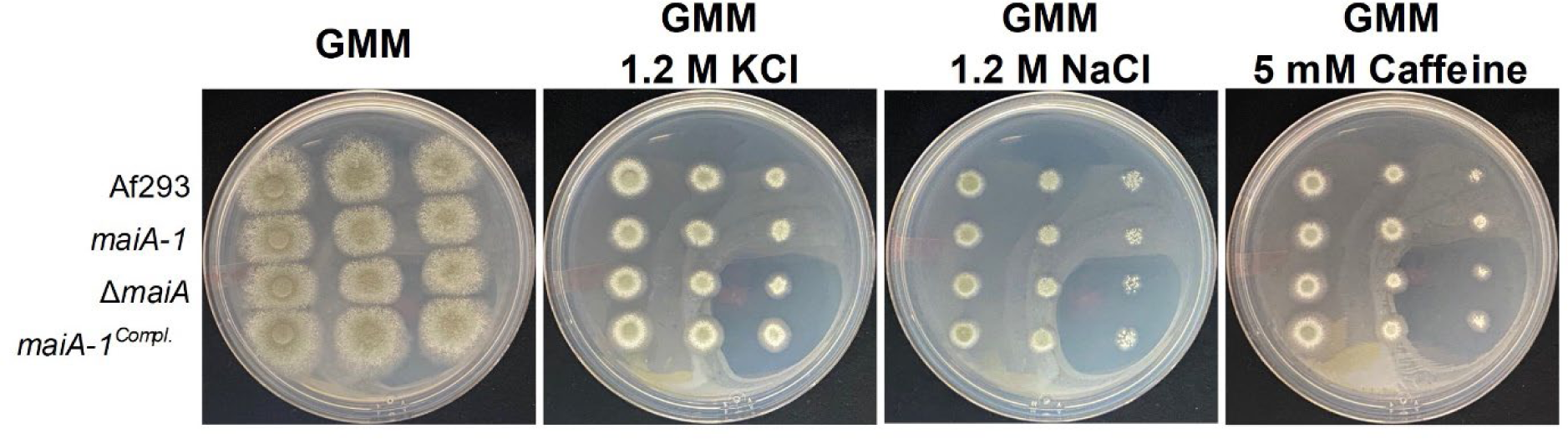
Phenotypic spotting assay characterization of Af293, *maiA-1*, Δ*maiA*, and *maiA-1^comp.^* using GMM agar plates supplemented with the indicated agents after 72 hours of incubation.

**Figure S2.**
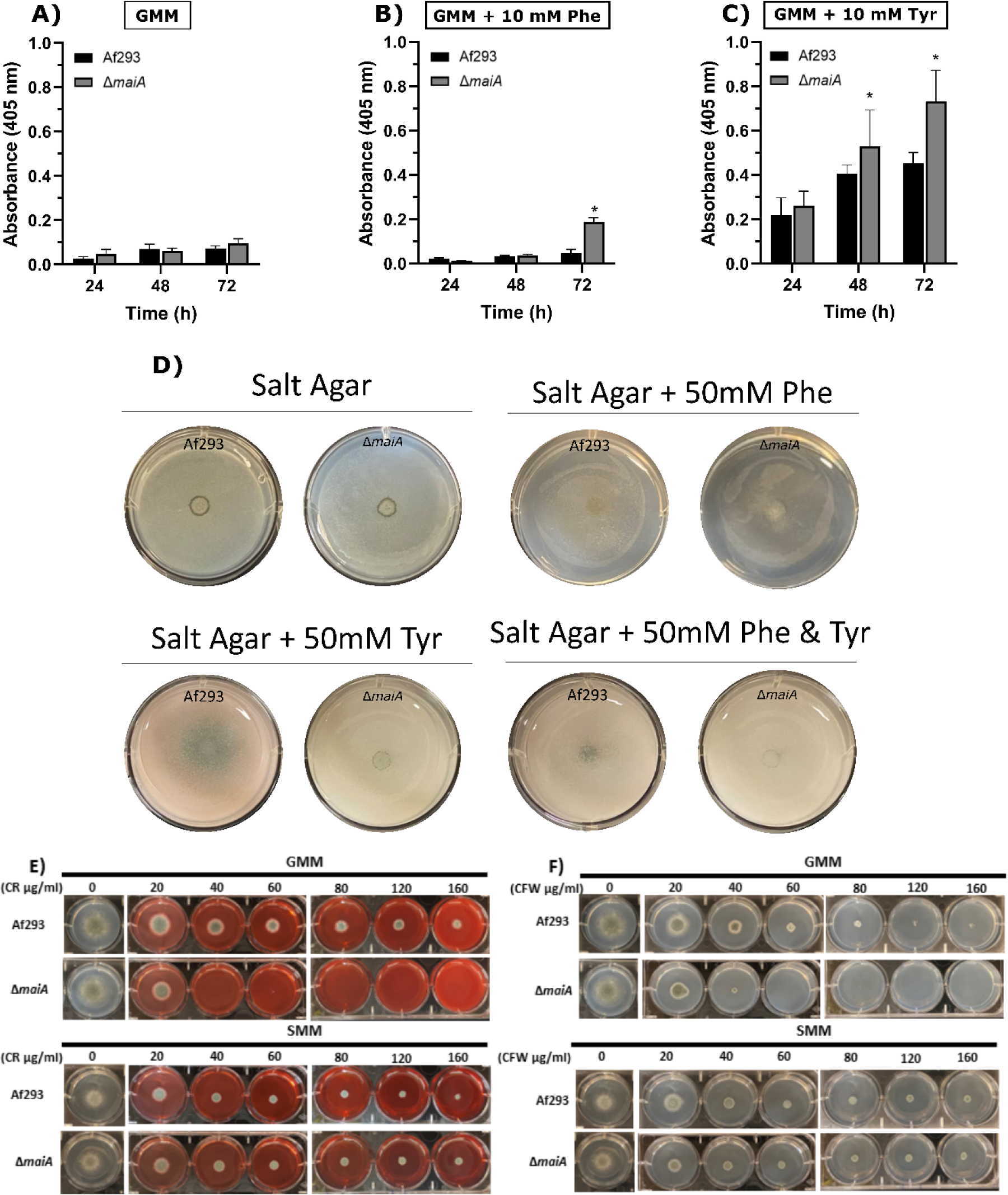
Phenotypic characterization of Δ*maiA* in response to Phe and Tyr to demonstrate that the phenotype of whole deletion of *maiA* gene is the same as the disruption strain *maiA-1*. Pyomelanin secretion ability of the Af293 and Δ*maiA-1* growing in **A)** GMM broth, **B)** GMM broth supplemented with 10 mM Phe and **C)** GMM broth supplemented with 10 mM Tyr. **D)** Spotting assay after 72 hours of incubation of the Af293 and Δ*maiA* strains on salt agar plates in which the only carbon source were Phe (50 mM), Tyr (50 mM) or Phe and Tyr (50 mM each). GMM or SMM 6-well plates supplemented with different concentrations (0, 20, 40, 60, 80, 120, 160 µg/ml) of congo red (CR) (**E**) or calcofluor white (CFW) (**F**).

**Figure S3.**
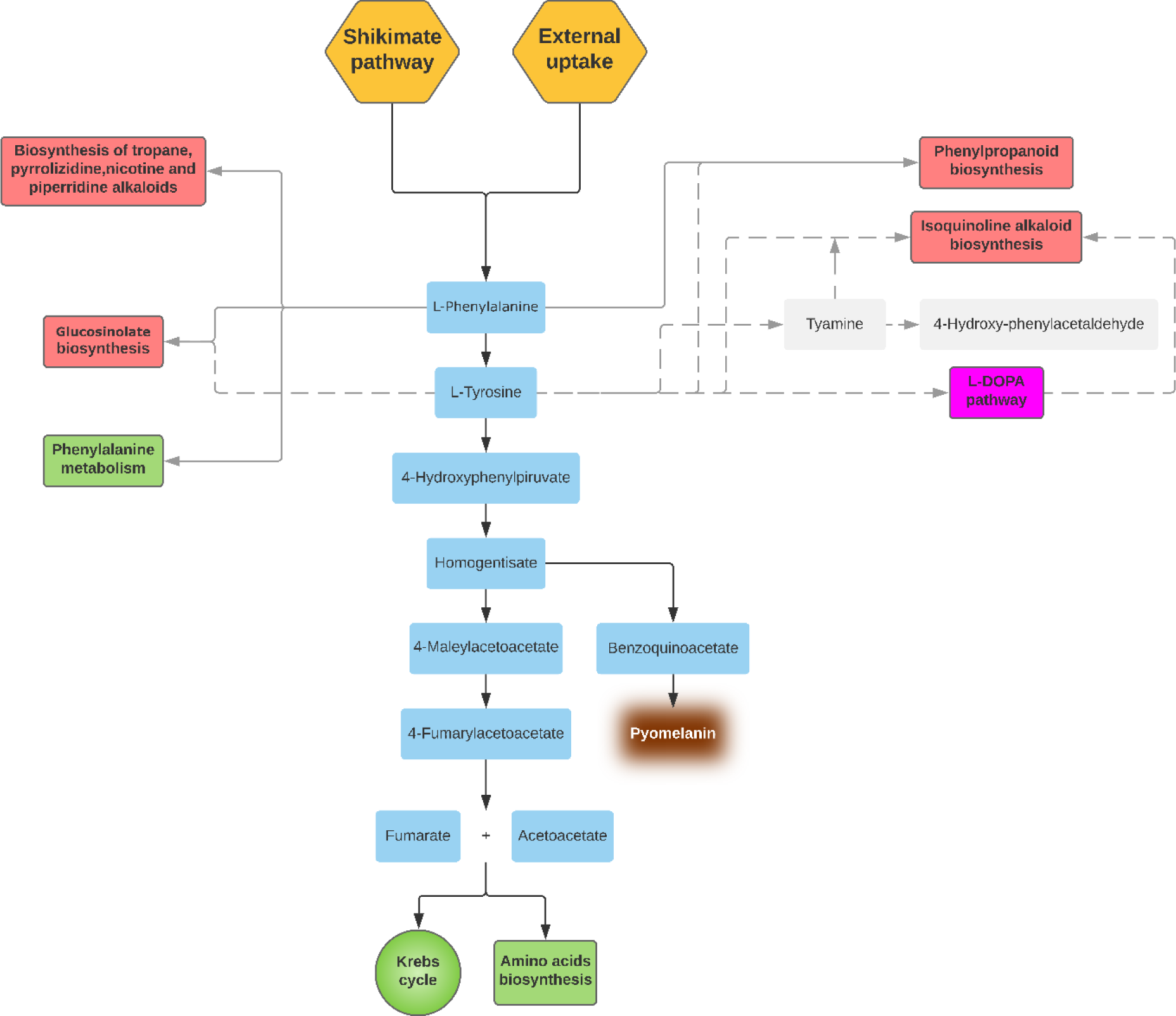
Schematic representation of the Phe/Tyr degradation pathway as well as the metabolic pathways in which Phe or Tyr take part. Red boxes indicate metabolic pathways still un-described in *A. fumigatus*. The pink box indicates that the pathway is described in *A. fumigatus* but do not generate energy as consequence. Green boxes represent metabolic pathways by which *A. fumigatus* could obtain energy. Orange boxes represent the two mechanisms of *A. fumigatus* to obtain Phe. Grey boxes represent steps described in other species but not in *A. fumigatus*. Grey boxes and dashed grey lines are related to Tyr while continuous grey lines are related to Phe. This figure was built in based on KEGG pathway database (https://www.genome.jp/kegg/pathway.html).

**Table S1.** Go enrichment analysis of the *A. fumigatus* up-regulated genes (FC > 1.5) and down-regulated genes (FC < −1.5) in both *in vitro* experimental models of infection.

| GO Slim Term | Number of Genes |  |  |  |
| --- | --- | --- | --- | --- |
|  | RAW 264.7 vs Control |  | A549 vs Control |  |
|  | Up-regulated | Down-regulated | Up-regulated | Down-regulated |
| **Biological process** | **126** | **101** | **403** | **189** |
| Transport | 35 | 34 | 89 | 40 |
| Regulation of biological process | 22 | 8 | 56 | 22 |
| Response to stress | 17 | 15 | 36 | 8 |
| Secondary metabolic process | 17 | 23 | 30 | 6 |
| Response to chemical | 14 | 17 | 40 | 11 |
| Lipid metabolic process | 13 | 8 | 30 | 3 |
| RNA metabolic process | 13 | 7 | 18 | 10 |
| Developmental process | 13 | 1 | 20 | 5 |
| Cellular amino acid metabolic process | 12 | 7 | 19 | 8 |
| Transcription, DNA-templated | 11 | 3 | 15 | 4 |
| Cellular respiration | 11 | - | 6 | 1 |
| Carbohydrate metabolic process | 10 | 20 | 49 | 23 |
| Toxin metabolic process | 9 | 2 | 12 | - |
| Cellular homeostasis | 7 | 18 | 9 | 7 |
| Cell cycle | 6 | 2 | 12 | 6 |
| Filamentous growth | 6 | 1 | 18 | 7 |
| Sexual sporulation | 6 | 1 | 7 | 2 |
| Asexual sporulation | 4 | - | 5 | 3 |
| Organelle organization | 3 | 1 | 16 | 11 |
| Pathogenesis | 3 | 7 | 12 | 4 |
| Translation | 2 | - | 1 | 2 |
| Cellular protein modification process | 2 | - | 9 | 10 |
| Conjugation | 1 | - | 3 | - |
| DNA metabolic process | 1 | - | 2 | 2 |
| Vitamin metabolic process | 1 | 2 | 1 | 2 |
| Cell adhesion | 1 | - | 3 | - |
| Signal transduction | 1 | - | 6 | 5 |
| Transposition | 1 | - | - | - |
| Ribosome biogenesis | - | 2 | 1 | 2 |
| Cytokinesis | - | 1 | 1 | 3 |
| Protein catabolic process | - | 1 | 2 | 4 |
| Cytoskeleton organization | - | - | 5 | 2 |
| Vesicle-mediated transport | - | - | 4 | 3 |
| Protein folding | - | - | 2 | - |
| Other | 35 | 34 | 105 | 52 |
|  | RAW 264.7 vs Control |  | A549 vs Control |  |
|  | Up-regulated | Down-regulated | Up-regulated | Down-regulated |
| **Cellular component** | **161** | **160** | **536** | **259** |
| Membrane | 55 | 33 | 112 | 49 |
| Mitochondrion | 32 | 4 | 28 | 5 |
| Nucleus | 27 | 8 | 37 | 18 |
| Cytosol | 20 | 3 | 10 | 5 |
| Plasma membrane | 17 | 8 | 32 | 7 |
| Extracellular region | 13 | 12 | 38 | 16 |
| Vacuole | 8 | 3 | 8 | 3 |
| Endomembrane system | 7 | 3 | 10 | 12 |
| Cell wall | 6 | 1 | 11 | 1 |
| Peroxisome | 6 | 3 | 9 | - |
| Endoplasmic reticulum | 5 | - | 2 | 6 |
| Golgi apparatus | 3 | - | 3 | 2 |
| Site of polarized growth | 1 | - | 7 | 3 |
| Nucleolus | - | 1 | - | 2 |
| Cell cortex | - | - | 6 | 1 |
| Cytoskeleton | - | - | 4 | 3 |
| Chromosome | - | - | 2 | 5 |
| Actin cytoskeleton | - | - | 1 | 1 |
| Microtubule cytoskeleton | - | - | 1 | 2 |
| Ribosome | - | - | - | 2 |
| Other | 11 | 11 | 41 | 9 |
|  | RAW 264.7 vs Control |  | A549 vs Control |  |
|  | Up-regulated | Down-regulated | Up-regulated | Down-regulated |
| **Molecular function** | **146** | **106** | **407** | **198** |
| Oxidoreductase activity | 62 | 37 | 117 | 54 |
| Hydrolase activity | 24 | 32 | 89 | 40 |
| Transporter activity | 21 | 18 | 48 | 13 |
| Transferase activity | 21 | 20 | 47 | 28 |
| DNA binding | 8 | 7 | 21 | 8 |
| Lyase activity | 8 | 2 | 14 | 5 |
| Peptidase activity | 7 | 1 | 10 | 5 |
| Protein binding | 3 | - | 8 | 2 |
| RNA binding | 2 | - | 5 | 4 |
| Isomerase activity | 2 | - | 5 | - |
| Protein kinase activity | 1 | - | 5 | - |
| Signal transducer activity | 1 | - | - | - |
| Structural molecule activity | 1 | - | - | - |
| Lipase activity | 1 | 1 | 4 | - |
| Phosphatase activity | 1 | - | 10 | 1 |
| Ligase activity | 1 | - | - | - |
| Enzyme regulator activity | 1 | 1 | - | - |
| Helicase activity | - | 1 | 1 | 2 |
| Ligase activity | - | - | 3 | 2 |
| Enzyme regulator activity | - | - | 3 | - |
| Nucleotidyltransferase activity | - | - | 1 | 1 |
| Structural molecule activity | - | - | 1 | 1 |
| Translation regulator activity | - | - | - | 1 |
| Other | 17 | 20 | 54 | 16 |

**Table S2.** List of common *A. fumigatus* up-regulated DEGs during the co-incubation of the fungus with the macrophages RAW 264.7 and the human lung epithelium cell line A549.

| ID | Product |  | Fold Change (log<sub>2</sub>) |  |
| --- | --- | --- | --- | --- |
|  |  |  | RAW 264.7 vs Control | A549 vs Control |
| Afu1g01300 | GPI anchored protein |  | 4.663 | 2.609 |
| Afu1g03570 | Acid phosphatase PHOa | *phoA* | 2.228 | 7.182 |
| Afu1g04160 | Aspartate aminotransferase |  | 3.223 | 2.452 |
| Afu1g06610 | NADH-quinone oxidoreductase, 23 kDa subunit |  | 2.774 | 3.180 |
| Afu1g07380 | Glutamate synthase Glt1 |  | 2.110 | 1.544 |
| Afu1g13510 | C6 transcription factor FacB/Cat8 |  | 2.392 | 2.545 |
| Afu1g13750 | C<sub>2</sub>H<sub>2</sub> transcription factor (Rpn4) |  | 1.663 | 2.095 |
| Afu1g14540 | Oxidoreductase, short-chain dehydrogenase/reductase family |  | 2.474 | 3.855 |
| Afu1g14550 | Mn superoxide dismutase MnSOD |  | 4.439 | 5.605 |
| Afu1g15590 | Succinate dehydrogenase subunit CybS |  | 2.927 | 1.708 |
| Afu1g17070 | FYVE domain protein |  | 2.640 | 3.023 |
| Afu1g17360 | bZIP transcription factor (BACH2) |  | 2.437 | 5.423 |
| Afu2g00630 | GDLS lipase/acylhydrolase family protein |  | 1.751 | 2.509 |
| Afu2g00890 | Conserved hypothetical protein |  | 1.553 | 2.057 |
| Afu2g02180 | Hypothetical protein |  | 1.565 | 1.343 |
| Afu2g02460 | Hypothetical protein |  | 2.462 | 4.392 |
| Afu2g03730 | Ctr copper transporter family protein |  | 2.334 | 3.116 |
| Afu2g04200 | 4-hydroxyphenylpyruvate dioxygenase |  | 6.384 | 6.783 |
| Afu2g04210 | Hypothetical protein |  | 3.843 | 4.234 |
| Afu2g04220 | Homogentisate 1,2-dioxygenase | *hmgA* | 3.782 | 3.726 |
| Afu2g04240 | Maleylacetoacetate isomerase | *maiA* | 2.416 | 2.219 |
| Afu2g04262 | C6 transcription factor |  | 2.331 | 1.539 |
| Afu2g05060 | Alternative oxidase AlxA |  | 4.657 | 5.252 |
| Afu2g07720 | Cytochrome b5 |  | 3.462 | 2.262 |
| Afu2g07910 | Myo-inositol transporter |  | 1.840 | 2.280 |
| Afu2g09400 | Cyclohexanone monooxygenase |  | 1.643 | 2.002 |
| Afu2g10540 | Hypothetical protein |  | 1.602 | 2.300 |
| Afu2g12630 | Allergenic cerato-platanin Asp f 13 | *aspf13* | 1.817 | 1.429 |
| Afu2g12710 | MFS monocarboxylate transporter. putative |  | 4.852 | 2.753 |
| Afu2g12780 | von Willebrand domain protein |  | 1.645 | 3.149 |
| Afu2g13175 | Hypothetical protein |  | 2.093 | 3.159 |
| Afu2g15860 | TAM domain methyltransferase |  | 3.356 | 1.866 |
| Afu2g16180 | Hypothetical protein |  | 1.616 | 1.682 |
| Afu2g16930 | Succinate:fumarate antiporter (Acr1) |  | 3.965 | 4.153 |
| Afu2g17790 | Amino acid transporter |  | 1.924 | 5.138 |
| Afu3g01150 | GPI anchored cell wall protein, putative |  | 3.154 | 2.034 |
| Afu3g03040 | Hypothetical protein |  | 2.144 | 3.023 |
| Afu3g03060 | Cell wall protein PhiA |  | 4.723 | 4.309 |
| Afu3g03230 | bZIP transcription factor |  | 1.760 | 1.932 |
| Afu3g03280 | FAD binding monooxygenase |  | 3.064 | 2.326 |
| Afu3g03290 | Hypothetical protein |  | 2.595 | 3.634 |
| Afu3g03310 | RTA1 domain protein |  | 3.383 | 4.797 |
| Afu3g06730 | MFS sugar transporter |  | 1.696 | 3.086 |
| Afu3g07810 | Succinate dehydrogenase subunit | *sdh1* | 2.721 | 1.519 |
| Afu3g07990 | GABA permease GabA |  | 1.622 | 4.559 |
| Afu3g08210 | Hypothetical protein |  | 1.729 | 2.506 |
| Afu3g09220 | P450 family fatty acid hydroxylase |  | 1.936 | 2.563 |
| Afu3g12920 | Non-ribosomal peptide synthase GliP-like |  | 2.194 | 2.061 |
| Afu3g13640 | Extracellular serine-rich protein |  | 1.779 | 2.691 |
| Afu3g13660 | Ctr copper transporter family protein |  | 5.135 | 4.005 |
| Afu3g13670 | Siderochrome-iron transporter |  | 4.181 | 2.705 |
| Afu3g13680 | Hypothetical protein |  | 4.238 | 3.022 |
| Afu3g13690 | Pyoverdine/dityrosine biosynthesis family protein, putative |  | 3.119 | 1.698 |
| Afu3g13700 | Transferase family protein |  | 6.523 | 5.245 |
| Afu3g14665 | Hypothetical protein |  | 4.943 | 4.108 |
| Afu3g14940 | Hypothetical protein |  | 2.490 | 2.713 |
| Afu4g01270 | Integral membrane protein |  | 3.144 | 1.868 |
| Afu4g01290 | Endo-chitosanase, pseudogene |  | 3.606 | 3.776 |
| Afu4g01470 | C6 finger domain protein |  | 2.026 | 2.629 |
| Afu4g03240 | Cell wall serine-threonine-rich galactomannoprotein Mp1 |  | 2.800 | 3.524 |
| Afu4g03270 | Epoxide hydrolase |  | 1.535 | 2.714 |
| Afu4g03410 | Flavohemoprotein |  | 4.829 | 3.099 |
| Afu4g03920 | MFS drug transporter |  | 3.569 | 5.353 |
| Afu4g03930 | Cysteine synthase B |  | 4.722 | 6.870 |
| Afu4g03940 | Ferric-chelate reductase |  | 3.165 | 3.276 |
| Afu4g04190 | Hypothetical protein |  | 3.355 | 4.413 |
| Afu4g04530 | Short chain dehydrogenase/reductase (Ayr1) | *ayr1* | 1.535 | 2.467 |
| Afu4g08490 | acyl-CoA dehydrogenase |  | 2.373 | 2.396 |
| Afu4g09110 | Cytochrome c peroxidase Ccp1 |  | 4.674 | 3.321 |
| Afu4g09140 | L-ornithine aminotransferase Car2 |  | 3.120 | 1.683 |
| Afu4g09470 | Cytochrome P450 monooxygenase |  | 2.532 | 3.903 |
| Afu4g09560 | ZIP Zinc transporter |  | 1.695 | 1.373 |
| Afu4g09580 | Major allergen Asp f 2 | *aspf2* | 2.676 | 2.452 |
| Afu4g10610 | Stress responsive A/B barrel domain protein |  | 1.591 | 1.942 |
| Afu4g10690 | Iron-sulfur cluster assembly accessory protein Isa1, putative |  | 3.122 | 2.066 |
| Afu4g13510 | Isocitrate lyase AcuD |  | 4.034 | 4.707 |
| Afu4g13540 | Potassium uptake transporter |  | 1.955 | 2.238 |
| Afu4g13780 | Polyphenol monooxygenase |  | 2.289 | 4.003 |
| Afu5g00300 | Zinc-binding oxidoreductase |  | 3.549 | 3.817 |
| Afu5g00740 | Hypothetical protein |  | 1.908 | 2.496 |
| Afu5g01030 | Glyceraldehyde 3-phosphate dehydrogenase |  | 1.927 | 3.091 |
| Afu5g01200 | Carboxypeptidase S1 |  | 2.858 | 2.204 |
| Afu5g02320 | Conserved hypothetical protein |  | 2.068 | 3.266 |
| Afu5g02330 | Major allergen and cytotoxin Asp f 1 (MitF) | *aspf1* | 3.001 | 3.102 |
| Afu5g07480 | Hypothetical protein |  | 1.705 | 1.695 |
| Afu5g07500 | β-lactamase family protein | | 1.761 | 2.657 |
| Afu5g09330 | CipC-like antibiotic response protein |  | 2.636 | 4.687 |
| Afu5g10370 | Iron-sulfur protein subunit of succinate dehydrogenase | *sdh2* | 2.986 | 1.928 |
| Afu5g11260 | Siderophore transcription factor | *sreA* | 2.104 | 2.095 |
| Afu5g11290 | D-amino acid oxidase |  | 1.551 | 1.559 |
| Afu5g12840 | Hydroxyacylglutathione hydrolase |  | 2.209 | 2.194 |
| Afu5g13800 | Transcriptional regulator |  | 2.291 | 2.068 |
| Afu5g13810 | Transulfuration enzyme family protein |  | 2.494 | 3.435 |
| Afu5g14650 | RING finger protein |  | 1.662 | 3.628 |
| Afu6g00290 | Aminotransferase |  | 1.678 | 1.504 |
| Afu6g00430 | IgE-binding protein |  | 2.836 | 2.241 |
| Afu6g00680 | Conserved hypothetical protein |  | 2.285 | 2.038 |
| Afu6g00690 | Conserved hypothetical protein |  | 3.229 | 3.073 |
| Afu6g00720 | LysM domain protein |  | 1.989 | 2.779 |
| Afu6g00740 | Conserved hypothetical protein |  | 2.983 | 4.180 |
| Afu6g02210 | Cytochrome P450 monooxygenase |  | 2.459 | 3.678 |
| Afu6g02810 | Ctr copper transporter |  | 1.810 | 1.767 |
| Afu6g02820 | Metalloreductase |  | 1.760 | 1.903 |
| Afu6g03540 | Malate synthase | *acuE* | 2.848 | 3.706 |
| Afu6g03590 | Citrate synthase | *cit1* | 2.610 | 3.541 |
| Afu6g07710 | Mitochondrial dicarboxylate carrier protein |  | 1.858 | 3.376 |
| Afu6g07720 | Phosphoenolpyruvate carboxykinase | *acuF* | 3.707 | 3.663 |
| Afu6g07750 | MFS phospholipid transporter | *git1* | 2.105 | 2.506 |
| Afu6g09200 | Hypothetical protein |  | 2.230 | 1.594 |
| Afu6g10310 | Hypothetical protein |  | 2.313 | 3.249 |
| Afu6g12250 | Succinyl-CoA:3-ketoacid-coenzyme A transferase (ScoT), putative | *scoT* | 2.072 | 3.276 |
| Afu6g12930 | Mitochondrial aconitate hydratase |  | 2.077 | 2.090 |
| Afu6g13750 | Ferric-chelate reductase |  | 2.445 | 1.820 |
| Afu6g14010 | GPI anchored protein |  | 3.245 | 2.255 |
| Afu7g00990 | Transcriptional activator of ethanol catabolism AlcS |  | 4.067 | 4.992 |
| Afu7g01020 | Hypothetical protein |  | 2.092 | 1.221 |
| Afu7g01050 | Salicylate hydroxylase |  | 2.203 | 2.215 |
| Afu7g02010 | Indoleamine 2,3-dioxygenase family protein |  | 5.696 | 3.663 |
| Afu7g02070 | AIF-like mitochondrial oxidoreductase (Nfr1) |  | 2.957 | 3.208 |
| Afu7g05490 | Conserved hypothetical protein |  | 3.083 | 3.841 |
| Afu7g05500 | Glutathione S-transferase |  | 2.045 | 1.817 |
| Afu7g06180 | Hypothetical protein |  | 2.220 | 3.612 |
| Afu7g06380 | Maltase |  | 1.663 | 1.977 |
| Afu7g06680 | AAA family ATPase |  | 1.735 | 2.200 |
| Afu7g06820 | Galactose oxidase |  | 1.519 | 2.193 |
| Afu7g07020 | Hypothetical protein |  | 2.404 | 4.156 |
| Afu7g07060 | Hypothetical protein |  | 2.285 | 4.371 |
| Afu7g08310 | Hypothetical protein |  | 2.245 | 4.523 |
| Afu8g00790 | Hypothetical protein |  | 3.704 | 2.542 |
| Afu8g01310 | Ferric-chelate reductase | *fre2* | 6.239 | 6.378 |
| Afu8g01670 | Bifunctional catalase-peroxidase | *cat2* | 1.828 | 2.289 |
| Afu8g02050 | Conserved hypothetical protein |  | 2.061 | 3.535 |
| Afu8g02060 | Glycan biosynthesis protein (PigL), putative |  | 3.115 | 4.641 |
| Afu8g02070 | Glycosyl transferase |  | 3.893 | 5.660 |
| Afu8g02090 | Nucleotide-sugar transporter |  | 2.299 | 3.432 |
| Afu8g02620 | CobW domain protein |  | 2.871 | 6.542 |
| Afu8g05530 | Fumarate reductase | *osm1* | 2.522 | 2.555 |
| Afu8g05730 | β-glucosidase | | 2.191 | 3.195 |
| Afu8g06080 | Flavohemoprotein |  | 4.032 | 3.217 |

**Table S3.**
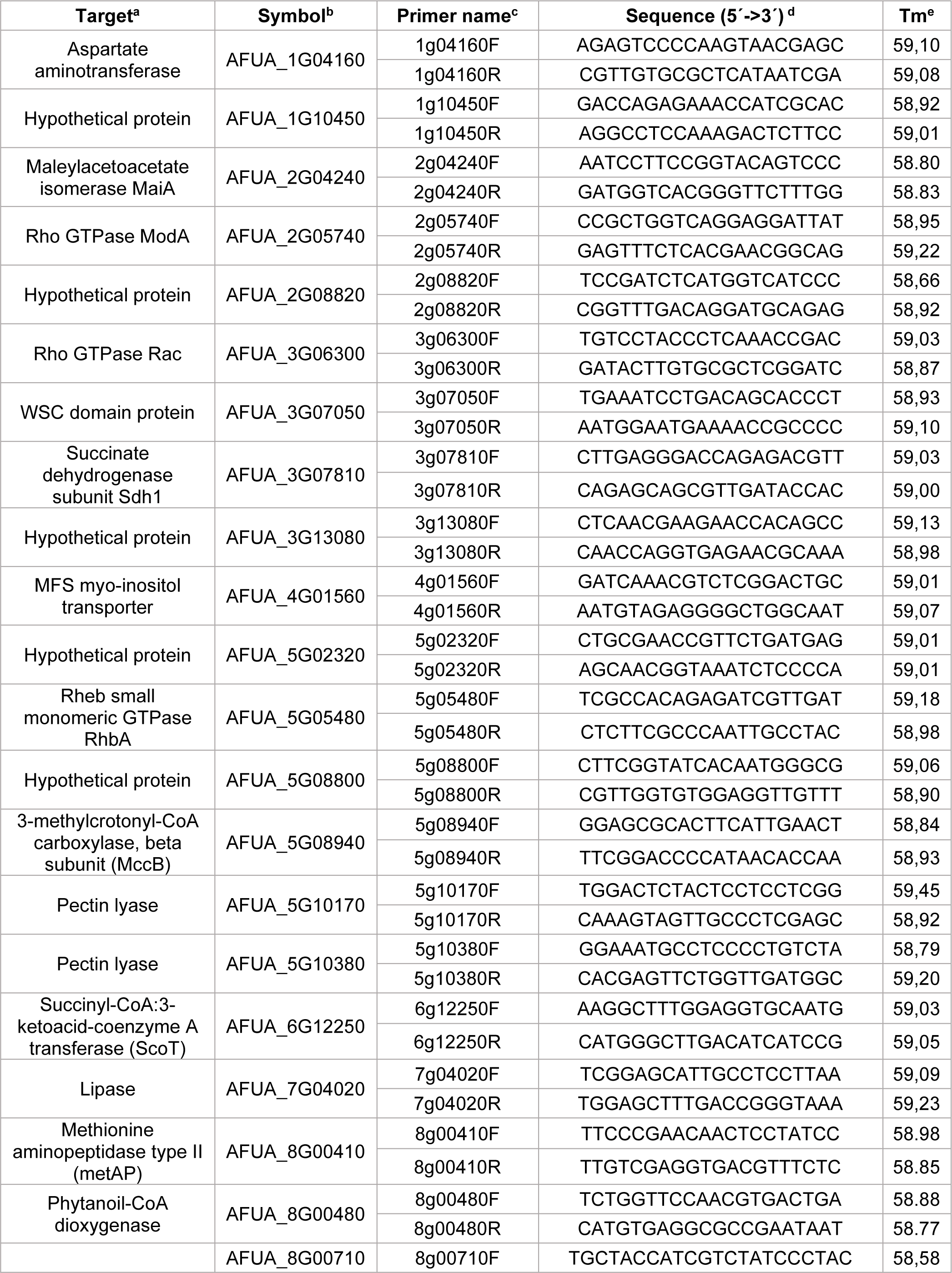
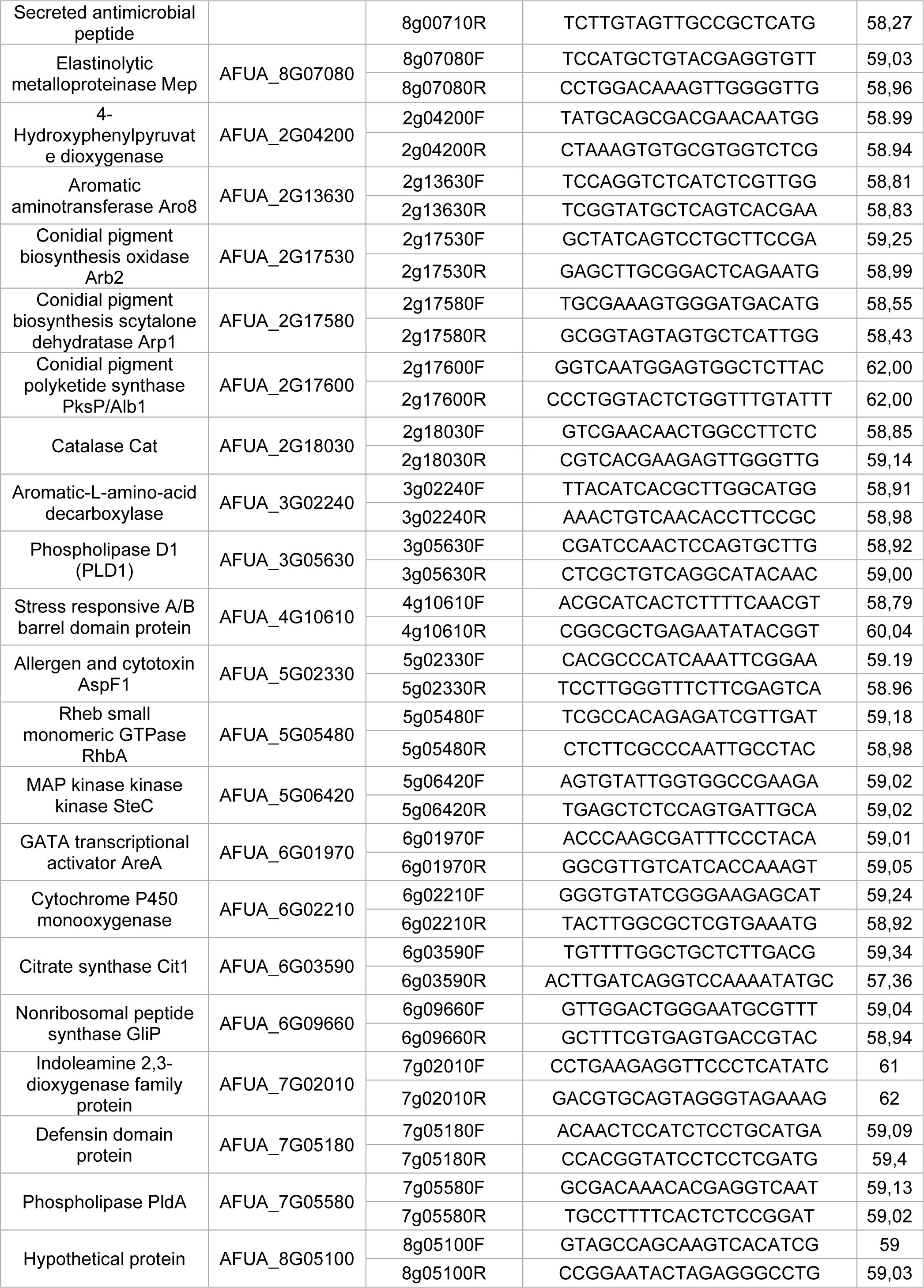
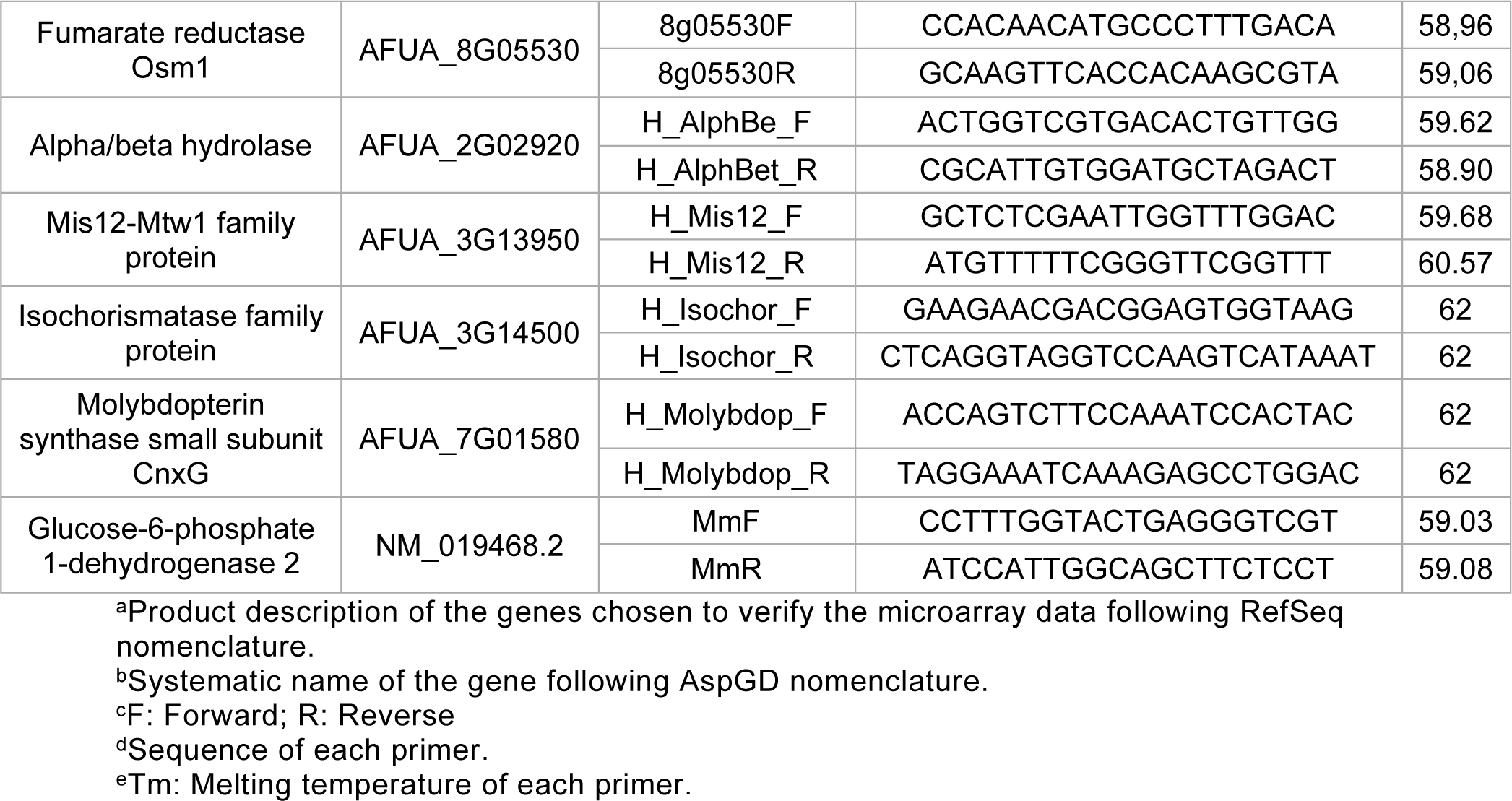
List of primers used for the microarray verification process.

